# Evolutionary changes in the chromatin landscape contribute to reorganization of a developmental gene network during rapid life history evolution in sea urchins

**DOI:** 10.1101/2022.02.22.481523

**Authors:** Phillip L. Davidson, Maria Byrne, Gregory A. Wray

## Abstract

Chromatin configuration is highly dynamic during embryonic development in animals, exerting an important point of control in transcriptional regulation. Yet there exists remarkably little information about the role of evolutionary changes in chromatin configuration to the evolution of gene expression and organismal traits. Genome-wide assays of chromatin configuration, coupled with whole-genome alignments, can help address this gap in knowledge in several ways. In this study we present a comparative analysis of regulatory element sequences and accessibility throughout embryogenesis in three sea urchin species with divergent life histories: a lecithotroph *Heliocidaris erythrogramma*, a closely related planktotroph *H. tuberculata*, and a distantly related planktotroph *Lytechinus variegatus*. We identified distinct epigenetic and mutational signatures of evolutionary modifications to the function of putative *cis*-regulatory elements in *H. erythrogramma* that have accumulated non-uniformly throughout the genome, suggesting selection, rather than drift, underlies many modifications associated with the derived life history. Specifically, regulatory elements composing the sea urchin developmental gene regulatory network are enriched for signatures of positive selection and accessibility changes which may function to alter binding affinity and access of developmental transcription factors to these sites. Furthermore, regulatory element changes often correlate with divergent expression patterns of genes involved in cell type specification, morphogenesis, and development of other derived traits, suggesting these evolutionary modifications have been consequential for phenotypic evolution in *H. erythrogramma*. Collectively, our results demonstrate that selective pressures imposed by changes in developmental life history rapidly reshape the *cis*-regulatory landscape of core developmental genes to generate novel traits and embryonic programs.

## INTRODUCTION

*Cis*-regulatory elements are significant contributors to phenotypic evolution and trait diversification by modulating transcriptional regulation via gene network interactions (Davidson and Erwin 2006; Wray 2007; Wittkopp and Kalay 2012). Extensive empirical evidence demonstrates that the function of *cis*-regulatory elements depends on distinct molecular characteristics such as the element’s sequence and epigenetic landscape that collectively provide a dynamic and heritable genomic substrate affecting inter- and intraspecific diversification. Enabled by advances in genomic technology and reduced sequencing costs, more recent work has demonstrated the importance of *cis*-regulatory elements in diverse biological contexts across a variety of animal systems at a genome-wide scale; for example: origination of human-specific traits (Shibata, et al. 2012; Swain-Lenz, et al. 2019), convergent evolution of flight loss in birds (Sackton, et al. 2019), population-level adaptation (Lewis and Reed 2019), genetic assimilation (van der Burg, et al. 2020) and color identity (Livraghi, et al. 2021) in butterfly wing patterning, embryonic regulation of gene expression in a crustacean (Sun and Patel 2021), limb bud development in chicken and mice (Jhanwar, et al. 2021), and single-cell resolution of *Drosophila* cell lineages (Cusanovich, et al. 2018). However, direct integration of large-scale *cis*-regulatory analysis and transcription in a defined evolutionary-developmental (evo-devo) context is lacking. In particular, relationships governing divergence in chromatin and gene expression, the role of such changes in trait evolution, and the action of natural selection on these processes remain largely unexplored within an interrogable study system providing resolved phylogenetic relationships, chromosome-level reference genomes, and sufficiently recent divergence times that noncoding regions can be aligned with confidence. In this study, we address these important questions by comparing the function of *cis*-regulatory elements in embryonic gene expression of sea urchin species with divergent developmental life history strategies to assess the extent *cis*-regulatory elements play in the evolution of gene expression, developmental systems, and by extension, trait variation and origination.

Among the most common and well-studied life history shifts in the animal kingdom is the evolution of feeding or non-feeding larval development (Flatt and Heyland 2011). In marine invertebrates, planktotrophy (feeding larval development) is characterized by species that produce many, relatively small eggs and have a longer planktonic larval period, whereas lecithotrophic species (non-feeding larval development) invest more maternal resources into a fewer number of large eggs that develop through an abbreviated larval stage (Strathmann 1985). Importantly, the ecological circumstances associated with each strategy are predictable and well-studied across diverse marine clades (Vance 1973; Strathmann 1985; Pechenik 1999; Marshall, et al. 2012; Morgan 2020), providing natural systems to analyze how selection shapes often rapid phenotypic divergence in developmental regulation and physiology. For example, within echinoderms, lecithotrophy has independently evolved from the ancestral planktotrophic state more than 20 times (Raff 1987; Wray 1996; McEdward and Miner 2001; Hart 2002), implying that recurring sets of environmental conditions (e.g. limited availability of phytoplankton) and selection can enact evolutionary modifications to even highly conserved developmental processes. The consequences include changes throughout development, ranging from egg composition (Prowse, et al. 2009; Falkner, et al. 2015; Byrne and Sewell 2019; Davidson, et al. 2019) and zygotic genome activation (Wray and Raff 1990; Israel, et al. 2016; Wang, et al. 2020), cell type specification and embryonic patterning (Wray and Raff 1989; Love and Raff 2006), to construction of larval structures such as the skeleton and imaginal adult rudiment (Emlet 1995; Koop, et al. 2017). Still, the impact selection exerts on the evolution of *cis*-regulatory interactions underlying alternative developmental programs and contrasting life history strategies is not known. This gap in knowledge is manifested more broadly in our current sparse understanding of the biological relationships governing regulatory element function, evolutionary constraints inherent to development and gene network architecture, and phenotypic diversification.

Perhaps the best studied example of the life history switch from planktotrophic to lecithotrophic development is the sea urchin *Heliocidaris erythrogramma*. Prior work has found dramatic modifications to morphogenesis and embryonic patterning in *H. erythrogramma* relative to planktotrophic sea urchin species (for reviews see Raff and Byrne 2006; Wray 2022). For instance, dorso-ventral and left-right axis establishment occurs considerably earlier in this species (Henry, et al. 1990), concordant with accelerated rudiment development and metamorphosis (Williams and Anderson 1975). At other embryonic domains, specification of cell lineages such as those responsible for mesenchymal tissue types have been delayed, lost, or spatially rearranged (Wray and Raff 1990; Klueg, et al. 1997; Wilson, Andrews, Rudolf Turner, et al. 2005; Davidson, et al. 2021). Recent transcriptomic studies have shown that expression profiles of key developmental gene regulatory network (dGRN) genes in this species have diverged from the highly conserved expression patterns in planktotrophic species (Israel, et al. 2016), and many of these regulatory changes are predicted to be based in *cis* (Wang, et al. 2020). However, the role of evolutionary changes in the sequence and configuration of *cis*-regulatory elements to the rapid evolution of these derived phenotype in *H. erythrogramma* is not known.

To address this, we compared chromatin accessibility, regulatory element sequence, and gene expression across six developmental stages in three sea urchin species spanning this life history switch: the lecithotroph *H. erythrogramma*, a closely-related (∼ 5 my divergence) planktotroph *H. tuberculata*, and a distantly-related (∼ 40 my divergence) planktotroph, *Lytechinus variegatus* (fig. 1a). Planktotrophic (feeding) larvae represent the *ancestral* life history mode within echinoderms relative to the *derived* lecithotrophic state (Strathmann 1978, Raff 1987, Wray 1996, McEdward and Miner 2001), so recent and dramatic changes in developmental phenotypes in *H. erythrogramma* provide an exceptional natural system to analyze the consequences of strong shifts in selection and life history on the evolution of gene regulatory mechanisms (Wray 2022).

**Figure 1:**
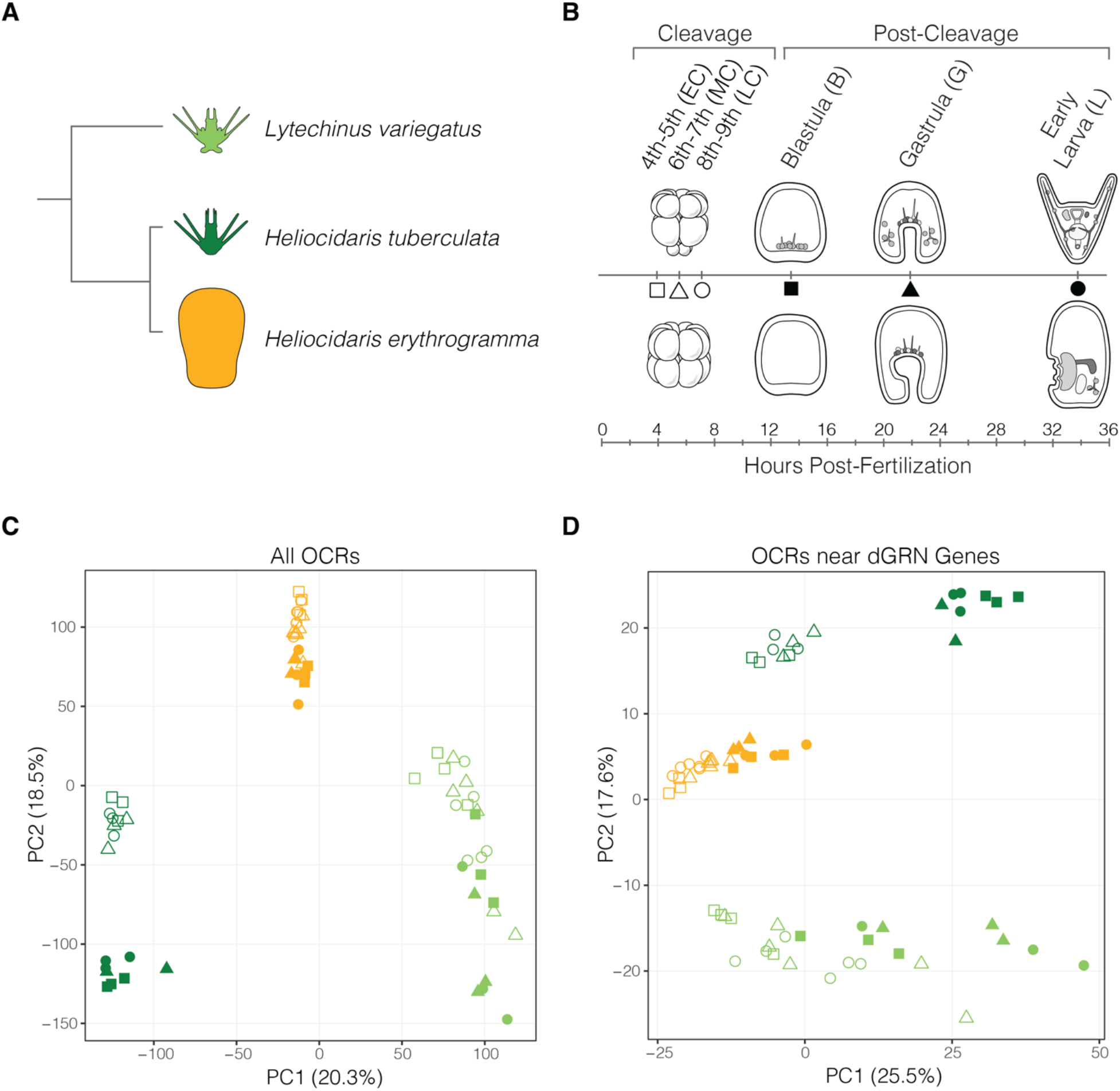
Global variation in chromatin accessibility between sea urchin embryos is largely driven by developmental stage and life history strategy. (*A*) Three focal sea urchin species of this study, including two planktotrophic species representing the ancestral life history (light green: *Lytechinus variegatus*; dark green: *Heliocidaris tuberculata*) and one lecithotrophic species representing the derived life history (orange: *Heliocidaris erythrogramma*). This color scheme is consistent throughout the study. (*B*) Developmental stages assayed in this study for each species. The ancestral life history is illustrated above, with corresponding developmental stages of the derived life history below. Developmental times are shown for *Heliocidaris tuberculata* at 21°C. Note, however, samples were collected on the basis of developmental stage, not hours post-fertilization, for all three species. (*C-D*) Principal component analysis of variation in normalized chromatin accessibility among (*C*) all orthologous OCRs and (*D*) only OCRs near dGRN genes through embryogenesis. OCR: open chromatin region; dGRN: developmental gene regulatory network.

In this study, we investigate the potential contribution of changes in regulatory element accessibility and sequence, as well as their interactions, on the evolution of developmental gene expression at a genome-wide scale. We consider our results in the context of two simple mechanistic models for an evolutionary change in the function of an individual regulatory element during development. (1) *local mutation first*: mutation that occurs within the regulatory element that alters the recruitment of a transcription factor, histone remodeler, or other local chromatin modifier (Clapier and Cairns, 2009; Zaret and Carroll, 2011), thereby altering the accessibility of the regulatory element and, in turn, possibly the transcription of a nearby gene. (2) *regulatory factor first*: a change elsewhere in the genome alters the expression of a pioneer transcription factor, histone remodeler, or other transcription factor that directly alters the accessibility of regulatory element in question and, in turn, possibly the transcription of a nearby gene. Both models accommodate an evolutionary increase or decrease in accessibility, and the accessibility change could alter interactions with other regulatory proteins, by masking or unmasking additional existing protein:DNA binding sites. Finally, in both models, the precipitating mutation(s) could be visible to selection through their impact on gene expression.

Within this framework, we identify distinct epigenetic and mutational signatures of evolutionary modifications in the noncoding genome of *H. erythrogramma* relative to species with the ancestral life history. Furthermore, we find that these changes have accumulated non-uniformly across the genome, and are notably enriched near genes with derived expression that are involved in cell type specification, morphogenesis, and development of other derived traits. The patterns of *cis*-regulatory modifications characterized here highlight the contribution of the epigenome to the evolution of transcriptional regulation and trait evolution more broadly.

## RESULTS AND DISCUSSION

We performed ATAC-seq on six developmental stages of *H. erythrogramma* (*He*), *H. tuberculata* (*Ht*), and *L. variegatus* (*Lv*) in order to identify open chromatin regions (OCRs) representing putative *cis*-regulatory elements (CREs) during embryogenesis. In this study, we use the term “OCR” when reporting analysis results because ATAC-seq is an assay of open chromatin. We apply the more specific term “CRE” when discussing the biological interpretation of our results because *cis*-regulatory elements are the operational units putatively identified within open chromatin peaks used to understand mechanisms of gene regulatory evolution. Importantly, however, while OCRs are known to be enriched for CRE sequences (Thurman et al., 2012), no strict relationship governs the presence of CREs based on occurrence of open chromatin peaks alone. Furthermore, even if an OCR is an authentic CRE, an evolutionary change in chromatin state may or may not have impact on the expression of a nearby gene. Indeed, many OCRs in sea urchin embryos open hours before any nearby gene is transcribed, including experimentally validated CREs (Shashikant et al. 2018; Arenas-Mena 2021); this finding suggests that the initial opening of many OCRs during early development makes a permissive rather than determinative contribution to activation of the zygotic transcriptome.

The developmental stages analyzed here include 4^th^/5^th^, 6^th^/7^th^, and 8^th^/9^th^ mixed cleavage stage embryos, as well as hatched blastula, mid-gastrula, and early larva (fig. 1b), enabling characterization of regulatory element dynamics from the onset of zygotic transcription through the specification and morphogenesis of complex tissue types. We then carried out alignments of chromosome-scale genome assemblies to find evolutionarily conserved OCRs between species to compare accessibility and sequence composition of putative CREs. These analyses provide a simultaneous comparison of gene regulatory mechanisms in two ways: (1) availability of CREs to transcription factor binding associated with differential chromatin configuration (accessibility) and (2) affinity of transcription factor binding to CREs via changes in nucleotide sequence, as evidenced by motif analysis and signatures of positive selection. Importantly, the genomes of these sea urchin species were assembled and annotated using an identical set of sequencing technology and methodology (Davidson, et al. 2020; Davidson, et al. 2021), minimizing technical bias and permitting high resolution identification of changes in regulatory element activity.

Of 42,679 OCRs identified across the three species, we found 35,788 1:1:1 orthologous, high confidence OCRs proximal to 11,679 gene models for cross-species comparison (fig. 2a). Note that inclusion in this set of 1:1:1 orthologues does not require that chromatin be open in all three species, but rather that it be open in at least one species at one time point surveyed and that the orthologous genomic sequence is present in all three species. This set of orthologous OCRs forms the basis of most analyses presented here. In addition to orthologous OCRs, we also identified sets of species-specific gains or losses of OCRs occurring on entire genomic segments present or lost in a single species (fig. 2a). Because these indel-based differences in OCRs are only present in at most two of the three sea urchin species compared in this study, they are treated as distinct sets of putative CREs. While the changes in regulatory element function examined in this study are not applicable to species-specific OCRs, it is important to note that many insertions containing species-specific OCR gains occur nearby genes with differential expression levels in *H. erythrogramma*, suggesting gain or loss of entire CREs may be an important component of the evolution of the derived life history strategy in this species (**supplementary fig. 1**).

**Figure 2:**
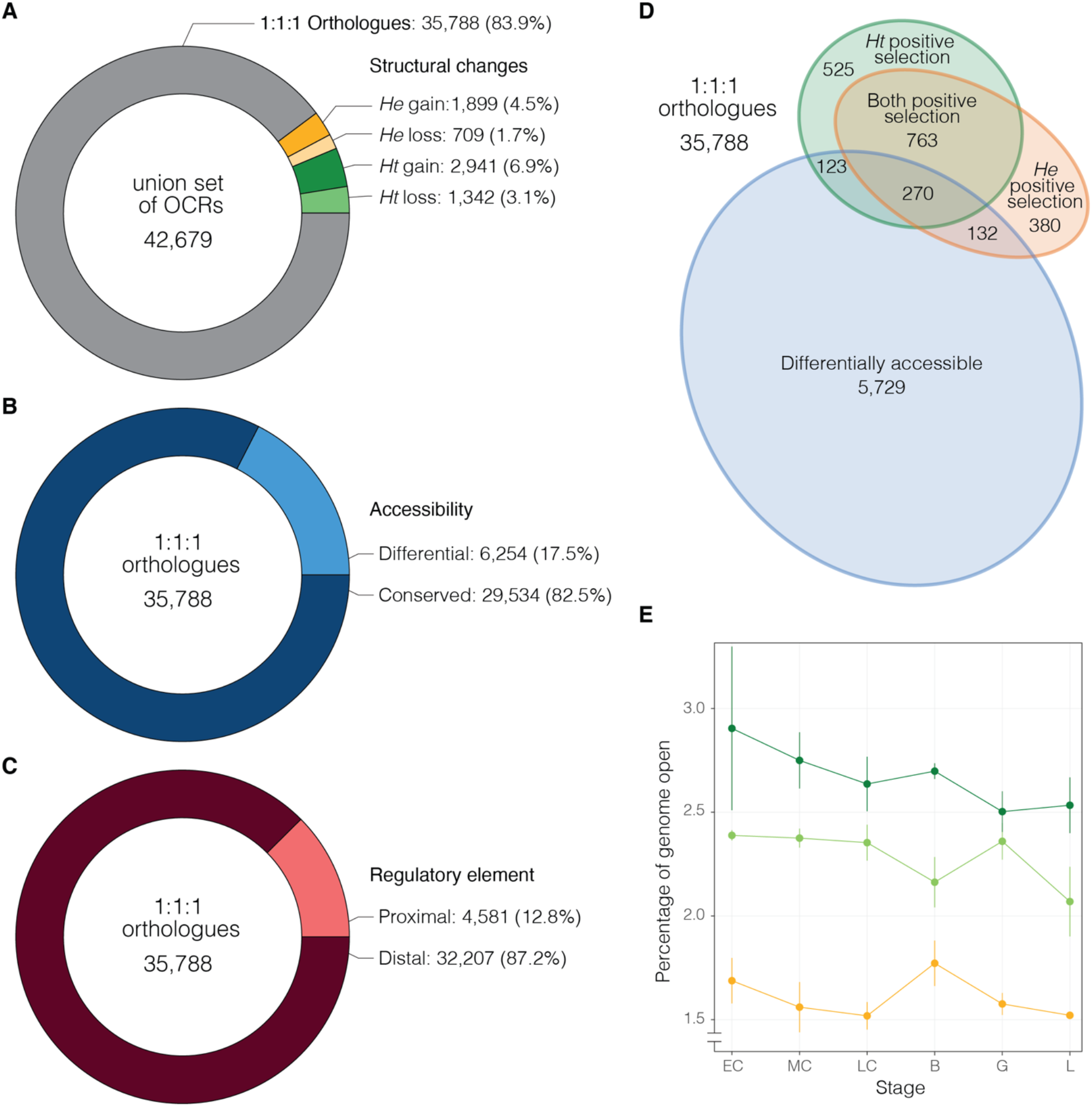
Summary statistics of regulatory element gains, losses, accessibility, and positive selection. (*A*) Union set of all OCRs, including structural differences between species such as lineage-specific gains and losses as well as 35,788 1:1:1 orthologous OCRs shared between all three species. These orthologous OCRs are the primary focus of this study. Breakdown of (*B*) OCR differentially accessibility between life history strategies and (*C*) OCR distance to gene models for these orthologous sites. (*D*) Overlap of OCR differential accessibility and evidence of positive selection on the *He*, *Ht*, or both branches. (*E*) Percentage of each species genome that is “open” at each stage of development. “Open” is defined as the cumulative portion of the genome represented by all OCRs (orthologous and lineage-specific) with significant levels of accessibility for a given stage and species (see **supplementary fig. 3** for additional information). Error bars indicate standard deviation of % openness across biological replicates for each species and developmental stage. OCR: open chromatin region; *He*: *Heliocidaris erythrogramma*; *Ht*: *Heliocidaris tuberculata*.

### Development and life history drive global variation patterns of accessibility

First, we assessed which biological factors may significantly contribute to variation in chromatin accessibility at a genomic scale. Principal component analysis (PCA) suggests species-level differences (PC 1: 20.3%) and the interaction of development and life history (PC 2: 18.5%) are the most important factors explaining variation in chromatin accessibility when all orthologous OCRs are considered (fig. 1c). Within each species, chromatin accessibility differences between stages roughly mirrors developmental progression as early and late stages cluster more closely to one another. Strikingly, though, there is far greater variation in accessibility through development in both species with the ancestral life history relative to *He*, suggesting the developmental chromatin landscape of the derived state is less dynamic during embryogenesis. This finding is consistent with delayed blastomere specification (Wray and Raff 1989) and simplified larval morphology in *He* (Emlet 1995), suggesting chromatin state changes normally associated with establishing cell identity and tissue differentiation have been modified during the evolution of its derived developmental life history.

Interestingly, when considering only OCRs near dGRN genes, the primary driver of variation in chromatin accessibility shifts to developmental life history rather than species-level differences (fig. 1d; PC 1: 25.5%), a finding corroborated by variance partitioning of the dataset (**supplementary fig. 2**). This result suggests chromatin configuration of critical developmental network genes is associated with developmental strategy in these species, as accessibility of regulatory elements controls the flow and deployment of embryonic programs, ultimately driving cell fate specification and morphogenesis. Therefore, changes in accessibility to CREs have the potential to modify the function and construction of novel developmental gene networks via altered regulatory interactions.

### Loss of regulatory element accessibility within the ancestral dGRN

We next quantified how the accessibility of 35,788 OCRs varies between life history strategies in order to better understand how chromatin configuration at specific CREs may influence the evolution and function of alternative embryonic programs. Across each cleavage stage examined (timepoints spanning 16-cell through ∼500-cell stage embryos), we found an average of 5.7% of OCRs were differentially accessible between ancestral and derived life histories, including a modest difference in sites more accessible in *Ht* and *Lv* (3.1%) or *He* (2.6%) (fig. 3a,b**; top**). Strikingly, starting at blastula stage, the proportion of differentially accessible OCRs across the genome rises, with most sites increasing in accessibility (“opening”) in the ancestral life history, but not in *He*. For example, by the larval stage, 9.5% of OCRs are differentially accessible between life histories, nearly three quarters of which (73.9%) are more accessible in *Ht* and *Lv*. Furthermore, when all OCRs are considered, including both orthologous and non-orthologous sites across species, a much greater proportion of the *He* genome is closed compared to *Ht* and *Lv*, suggesting that, at a genome-wide scale, *He* is characterized by zygotic quiescence during early development relative to ancestral life history (fig. 2e; **supplementary fig. 3**). Our results are consistent with the prediction that, as embryonic development proceeds, more regulatory sites open and become available to regulatory factor binding as zygotic transcription activates, cell lineages differentiate, tissue types are specified, and morphological structures are constructed. However, this genome “awakening” and reconfiguration of chromatin state across the embryo unfolds at a much grander scale in *Ht* and *Lv* relative to *He*. This finding supports the idea that embryonic fate specification in *He* is delayed relative to the ancestral state (Wray and Raff 1989, 1990) and collectively, is concordant with a morphologically simplified larva (Williams and Anderson 1975) and derived gene expression patterning in this species (Wilson, Andrews and Raff 2005; Wilson, Andrews, Rudolf Turner, et al. 2005; Smith, et al. 2008; Israel, et al. 2016; Koop, et al. 2017).

**Figure 3:**
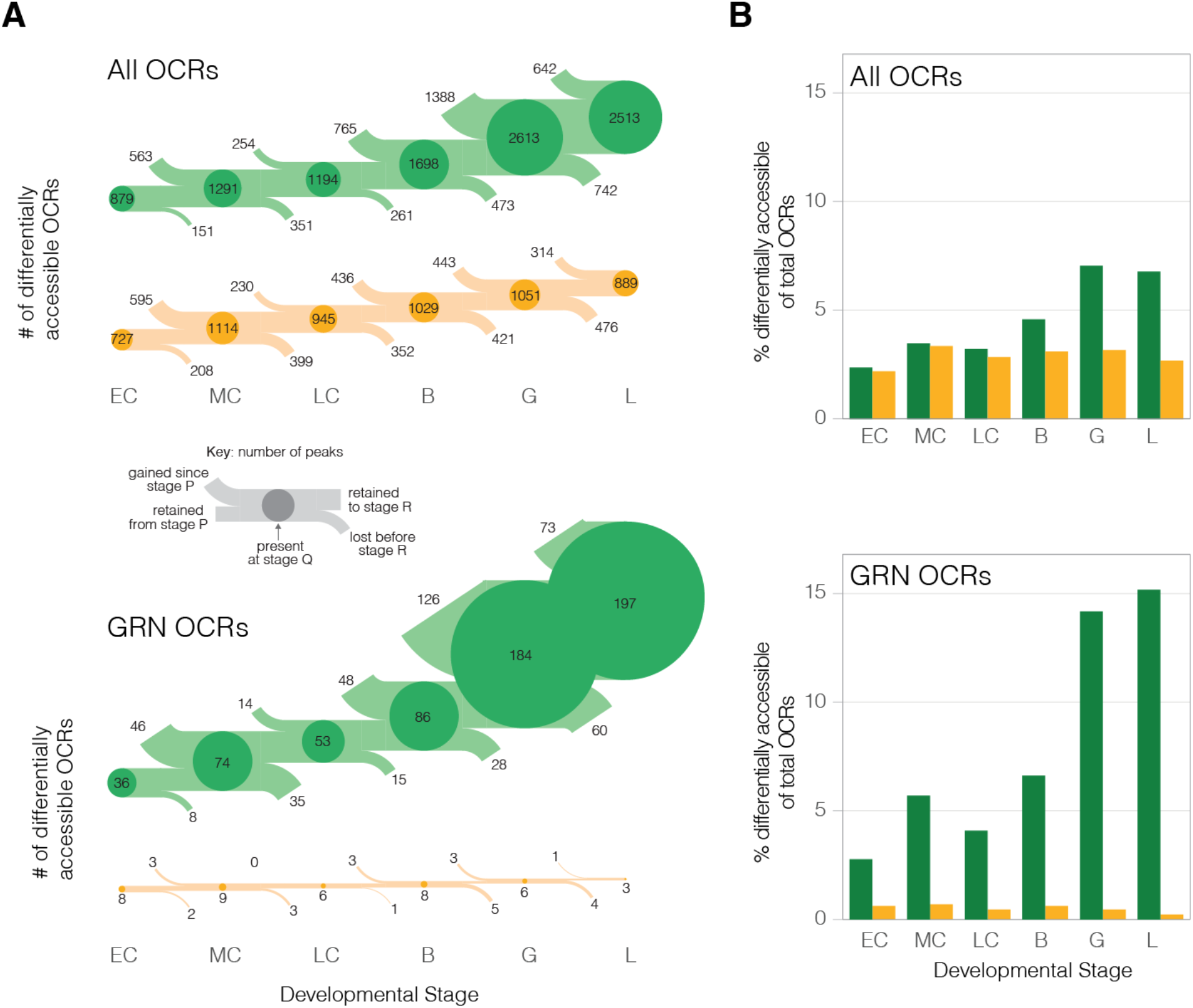
The chromatin landscape of the derived life history is far less accessible than that of species with the ancestral life history mode, particularly near core dGRN genes. (*A*) Number (flow diagram) and (*B*) percentage (barplots) of OCRs that are significantly differentially accessible in *H. erythrogramma* (orange) or species with the ancestral life history strategy (green) for all orthologous OCRs (top) and only OCRs near dGRN genes (bottom). If a differentially accessible OCR is more open in species with the ancestral developmental mode or in *H. erythrogramma*, it is included in the green or orange OCR sets, respectively. A key for flow diagrams is provided to explain how the number of differentially accessible OCRs are gained, retained, and lost between developmental stages. Circle size is proportional to the percentage of each set of OCRs (those near all genes or only dGRN genes) that are differentially accessible at a given stage. OCR: open chromatin region; dGRN: developmental gene regulatory network.

This delayed zygotic activation hypothesis is further supported when OCR accessibility of dGRN genes is compared. Near these critical developmental regulatory genes, we find a remarkable asymmetry in putative CRE accessibility between life history modes from the onset of development: on average, 0.5% of OCRs near dGRNs are more accessible in *He* during development, whereas this figure ranges from 2.8% (early cleavage) to 15.1% (larva) for species with the ancestral life history (fig. 3a.b**; bottom**). Therefore, modifications to CRE accessibility in *He* are not uniformly distributed across the genome, as would be expected of neutral processes such as drift, but instead are strikingly enriched near dGRN genes (Fisher’s exact test, two-sided: p < 1.31 x 10^-11^). An example of possible CRE activation in the ancestral state that is not found in *He* is the chromatin landscape proximal to the gene *patched* (*ptc*) (fig. 4). This gene encodes a receptor of the Hedgehog ligand; its misexpression results in skeletal defects and abnormal mesodermal patterning in the ancestral sea urchin developmental program (Walton, et al. 2009).

**Figure 4:**
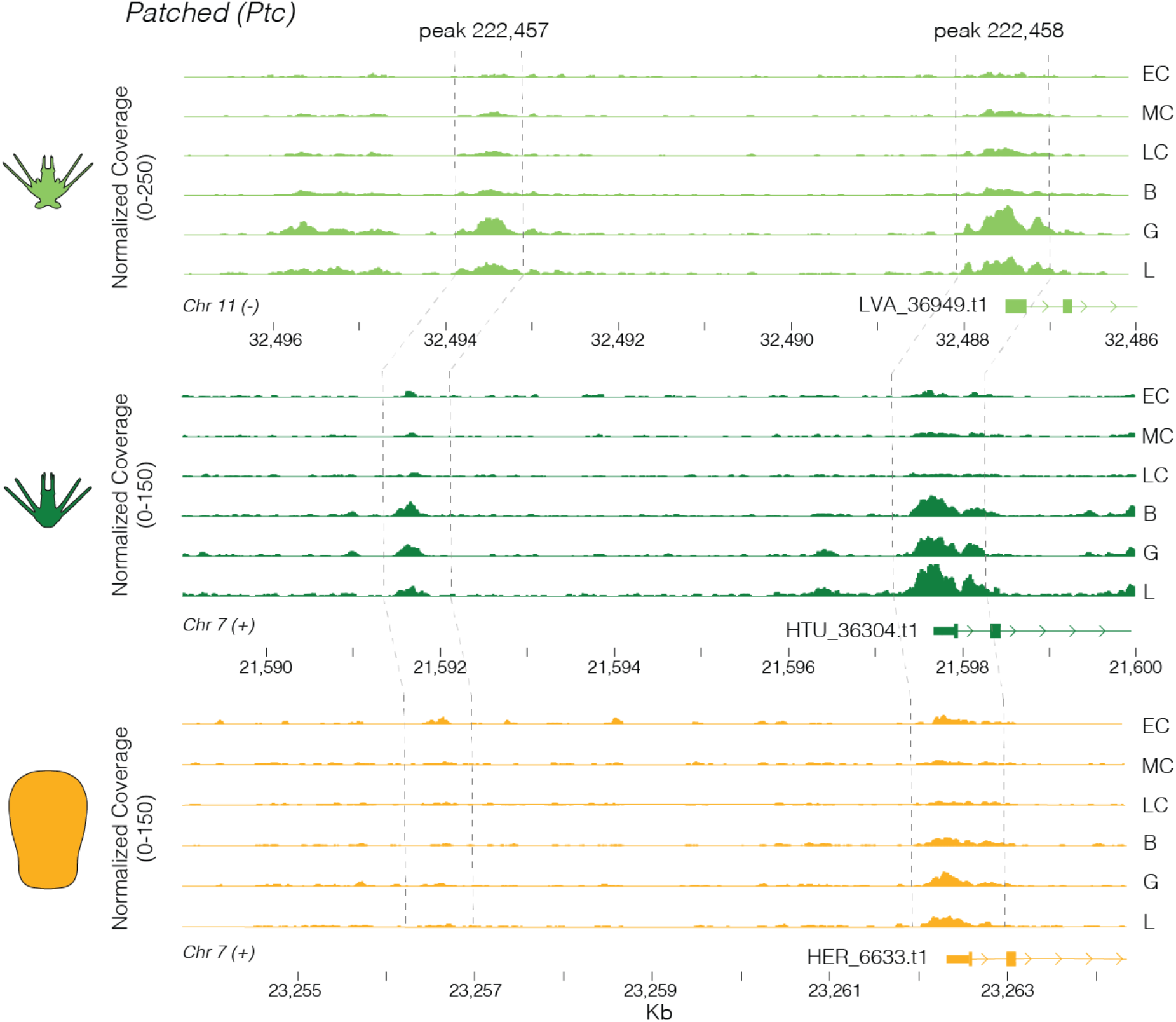
Example of an evolutionary change in chromatin accessibility during early development. The embryonic chromatin landscape near the gene *Patched* is shown for the three study species, with gene models on the right and dashed lines indicating orthologous noncoding regions. In the two planktotrophs, the most proximal OCR (likely promoter based on location at the 5’ end of the 5’ UTR) opens substantially after cleavage, and a distal OCR ∼7kb upstream opens by gastrula stage. In in *H. erythrogramma*, the proximal OCR is substantially less accessible and the distal OCR is not above background accessibility levels.

Given that certain ecological conditions likely favor the evolution of non-feeding development (Strathmann, 1978, 1985; Marshall, et al. 2012), the finding that differences in chromatin accessibility are concentrated around developmental regulatory genes suggests natural selection has sculpted the epigenetic landscape nearby the sea urchin dGRN during life history evolution, generating potentially adaptive changes to transcriptional regulation and expression patterning during embryogenesis. Furthermore, our results indicate that the evolutionary changes in putative CRE accessibility have functional consequences — if they had no impact on gene expression or some other trait, natural selection would not influence their status during development.

### Developmental regulatory elements have accumulated mutational signatures of positive selection

Prior studies have found enrichment of signals of positive selection in OCRs near genes that are differentially expressed between species (Edsall et al. 2019; Swain-Lenz et al. 2019; Shibata et al. 2021). To test whether a similar relationship exists in our study clade and whether it was accentuated during the evolution of lecithotrophy, we next analyzed sequence evolution of orthologous regulatory loci to identify instances of mutational pileup as a signal of positive selection within these sites. Substitution rates of 35,788 OCRs on the *He* and *Ht* branches (table S1) were compared to 10 randomly selected neutrally evolving, non-coding reference sequences, and a likelihood ratio test was implemented to fit substitution rates across the tree to a branch-specific positive selection model, using *Lv* as the outgroup species (see **Methods and Materials** for details). Our goal was to identify cases of gene regulatory evolution via nucleotide variation within CRE sequences, potentially acting to modify transcription factor binding affinity of these elements and associated changes to gene network expression and connectivity.

In total, we identified 648 OCRs with evidence of positive selection exclusively in the *Ht* genome, compared to 512 exclusively in the *He* genome (fig. 5a: green and orange points, respectively; see also fig. 2d). This result thus suggests that, while the life history of these two sea urchin species dramatically differs, many potential changes in gene regulation associated with other aspects of their biology have accumulated within each species. At a genome-wide scale, these putatively adaptive changes have occurred perhaps even more so in *Ht*, the species with ancestral developmental life history.

**Figure 5:**
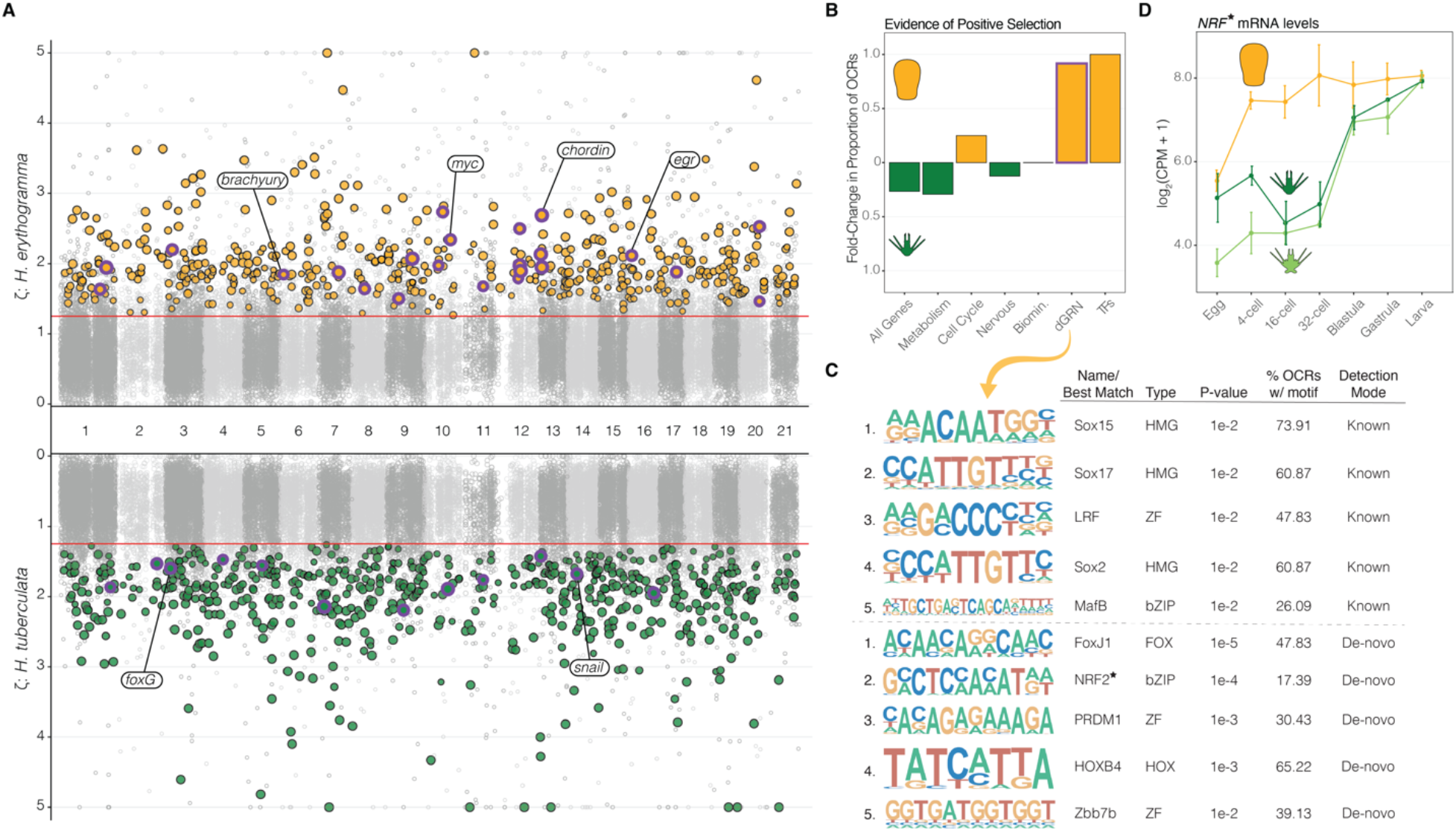
Sequence changes indicative of positive selection are enriched within *cis*-regulatory elements near dGRN genes and other transcription factors in the *H. erythrogramma* genome. (*A*) Zeta values (ratio of OCR substitution rate to that of neutrally evolving reference sequences) for *H. erythrogramma* (top) and *H. tuberculata* (bottom) across the genome. OCRs with significant support for lineage-specific evidence of positive selection are noted for *H. erythrogramma* (orange) and *H. tuberculata* (dark green), and those nearby dGRN genes are outlined in purple. (*B*) Fold-change in proportion of OCRs with more frequent evidence of positive selection in either *H. erythrogramma* or *H. tuberculata*, partitioned by functional categories of nearby genes (see **supplementary fig. 4** for all categories). (*C*) Transcription factor binding motif enrichment results of OCRs near dGRN genes with evidence of positive selection in *H. erythrogramma*. Detection mode indicates OCR sequences were searched against a (i) “known” set of optimized motif models or (ii) enriched motifs within the OCR were identified “de-novo” and matched to a known set of motifs. (*D*) Embryonic mRNA expression levels for *Nuclear factor erythroid 2-related factor* (*NRF*) for each species. OCR: open chromatin region; dGRN: developmental gene regulatory network.

Partitioning OCR selection results by the function of the nearest gene reveals only limited enrichment of positive selection within the putative regulatory elements of most functional classes of genes (fig. 5b, **supplementary fig. 4**). A prominent exception to this finding involves OCRs near genes in the sea urchin dGRN or near genes encoding other transcription factors. In these cases, approximately twice as many OCRs exhibit evidence of positive selection on the *He* branch relative to *Ht*. This result is especially striking considering *Ht* has 26.3% more OCRs with branch-specific evidence of positive selection across the entire genome (previous paragraph and fig. 5b). Moreover, nucleotide sequences of the 23 dGRN OCRs under selection in *He* are highly enriched for binding motifs of critical developmental transcription factors, including three SOX, three zinc finger, and one homeobox protein (fig. 5c). The SOX binding motifs are interesting because many transcription factors in this family act as “pioneer factors” that are capable of initiating chromatin opening (Zaret and Carroll 2011). Another interesting example involves NRF2: the binding motif for this transcription factor is overrepresented in OCRs under selection in *He* (fig. 5c) and the gene is expressed at much higher levels in *He* embryos relative to those in the ancestral life history (fig. 5d). This protein has gene regulatory functions in diverse biological contexts ranging from antioxidant response and cellular defense to metabolism and stem cell quiescence (Tonelli, et al. 2018), which in this case, could be related to derived embryonic detoxification and physiological systems (Tsushima, et al. 1995; Davidson, et al. 2019) or early cell lineage specification programs (Wray and Raff 1989, 1990; Davidson, et al. 2021) of *He*.

Taken together, putative regulatory elements tasked with wiring the ancestral sea urchin dGRN have accumulated a disproportionate number of mutations on the *He* branch. These likely play a significant role in regulation of divergent gene expression patterns and its highly derived developmental program. However, determining which mutations drive the signal of positive selection and their contribution to gained or lost transcription factor binding motifs remains a challenge. More broadly, our results provide strong evidence that natural selection can efficiently generate heritable changes concentrated within the *cis*-regulatory landscape of specific subsets of genes or gene networks to rapidly produce changes in transcriptional regulation.

### Overlap of evolutionary changes in regulatory element accessibility and sequence

The previous two sections describe how two distinct modifications to regulatory element function, namely accessibility and sequence, have accumulated across the *He* genome, and particularly so near critical developmental network genes. We next asked whether there exists any interaction between these two evolutionary changes to better understand the extent to which both epigenetic and sequence changes affect the same regulatory element. At all three post-cleavage stages, we found that differentially accessible OCRs contain a significantly greater proportion of sites with evidence of selection on the *He* branch relative to non-differentially accessible OCRs, including as much as an ∼50 % proportional increase at the blastula stage (Fisher’s exact test, two-sided: p < 5 x 10^-3^) (fig 6a: top). This same trend is observed at early cleavage (16/32-cell embryos), although not statistically significant. In contrast, we detected no statistical enrichment for positive selection within differentially accessible OCRs in the *Ht* genome; at early cleavage this relationship does approach significance, but no such trend exists at the later developmental stages (fig. 6a: bottom) (Fisher’s exact test, two-sided: p > 0.05).

**Figure 6:**
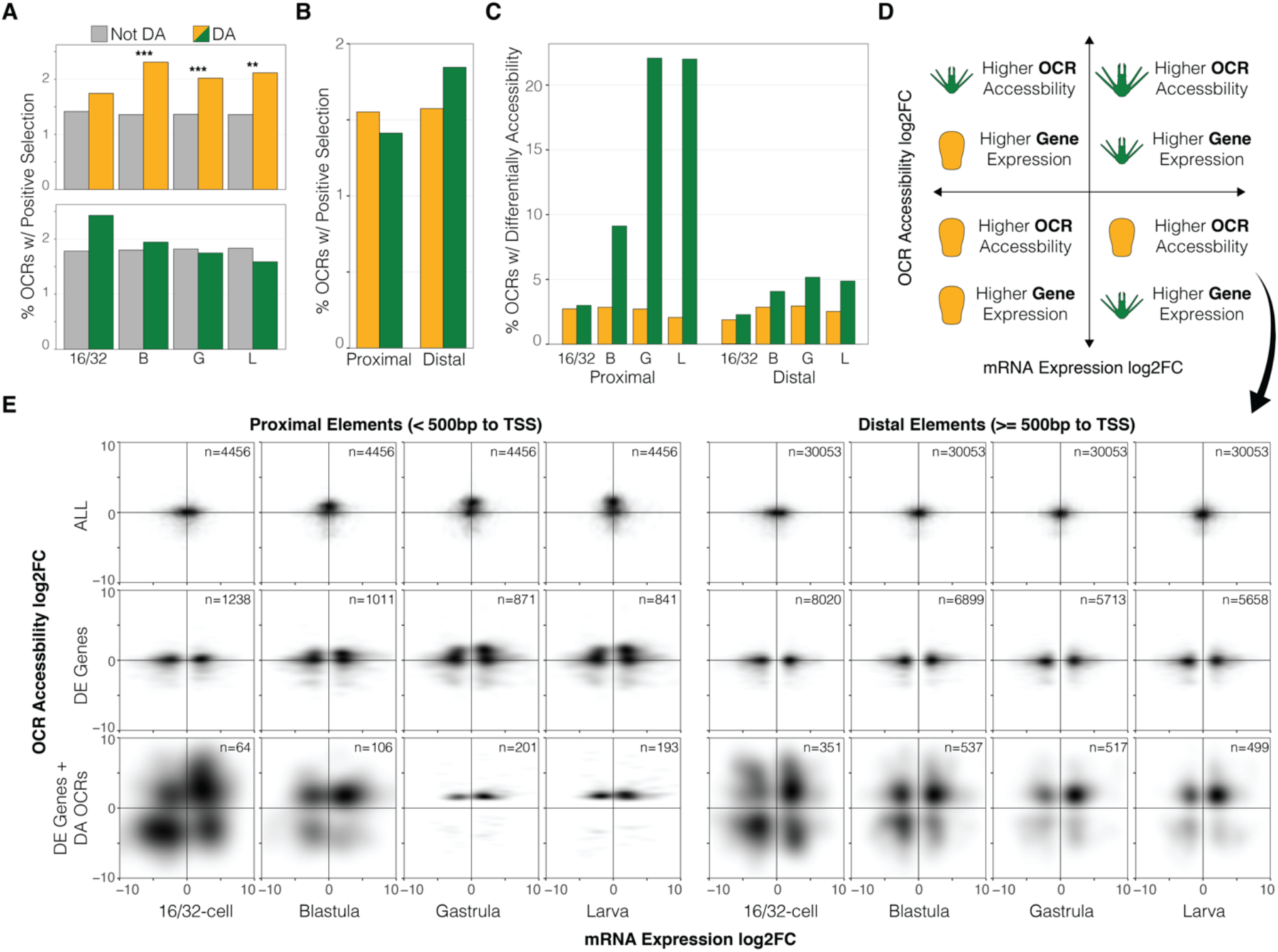
Independent and synergistic changes to regulatory element function have evolved during life history evolution in *H. erythrogramma*. (*A*) Proportion of differentially accessible or non-differentially accessible OCRs with evidence of positive selection on the *H. erythrogramma* branch (orange, top) or *H. tuberculata* branch (dark green, bottom). Two-sided, Fisher’s exact test: ** p ≤ 5 x 10^-3^; *** p ≤ 5 x 10^-4^. (*B*) Proportion of proximal of distal OCRs with evidence of positive selection in either *Heliocidaris* species. (*C*) Differential accessibility of proximal and distal OCRs at four stages of development. Orange indicates significantly greater accessibility in *H. erythrogramma*, dark green indicates significantly greater accessibility in *H. tuberculata*. (*D*) Diagram describing possible correlative patterns between changes in OCR accessibility and gene expression in either life history strategy (green: ancestral, planktotroph; orange: derived, lecithotroph): the layout of this panel is a key for interpreting the density plots in panel (*E*) below. (*E*) Fold-change of mRNA expression vs nearby OCR accessibility between life history strategies for proximal (left) and distal (right) OCRs. Relationships are shown for all genes and OCRs (*E*: top row), significantly differentially expressed genes (*E*: middle row), and significantly differentially expressed genes *and* differentially accessible OCRs (*E*: bottom row). OCR: open chromatin region; DE: differentially expressed (genes); DA: differentially accessible (OCRs).

Significant overlap in differential accessibility and positive selection in the *He* genome suggests a possible biological interaction between the two evolutionary mechanisms that may operate in tandem to alter regulatory function. It therefore becomes useful to consider the order in which these molecular events could take place (fig. 7). Measuring chromatin accessibility in hybrids provides a way to distinguish *cis* from *trans* genetic contributions to differences in individual OCRs (Connelly et. al 2014), using an approach originally developed for investigating the genetic basis for evolutionary changes in transcription (Wittkopp, et al. 2008; Crowley, et al. 2015). In the present case, the disproportionate occurrence of positive selection within differentially accessible OCRs only in the *He* genome suggests a potentially synergistic set of gene regulatory processes may operate to drive developmental change during the evolution of its derived life history. Because the tests for positive selection identify regions containing multiple derived mutations, additional *cis* mutations likely accumulated after the accessibility change, regardless of which molecular mechanism originally opened the region (fig. 7, t=3). The observation that this association is restricted to the *He* branch suggests that this interaction may often produce adaptive changes in transcription during development. More broadly, tandem accessibility and mutational changes may be a common gene regulatory method to efficiently modify gene expression for organismal adaptation and innovation across organismal lineages.

**Figure 7.**
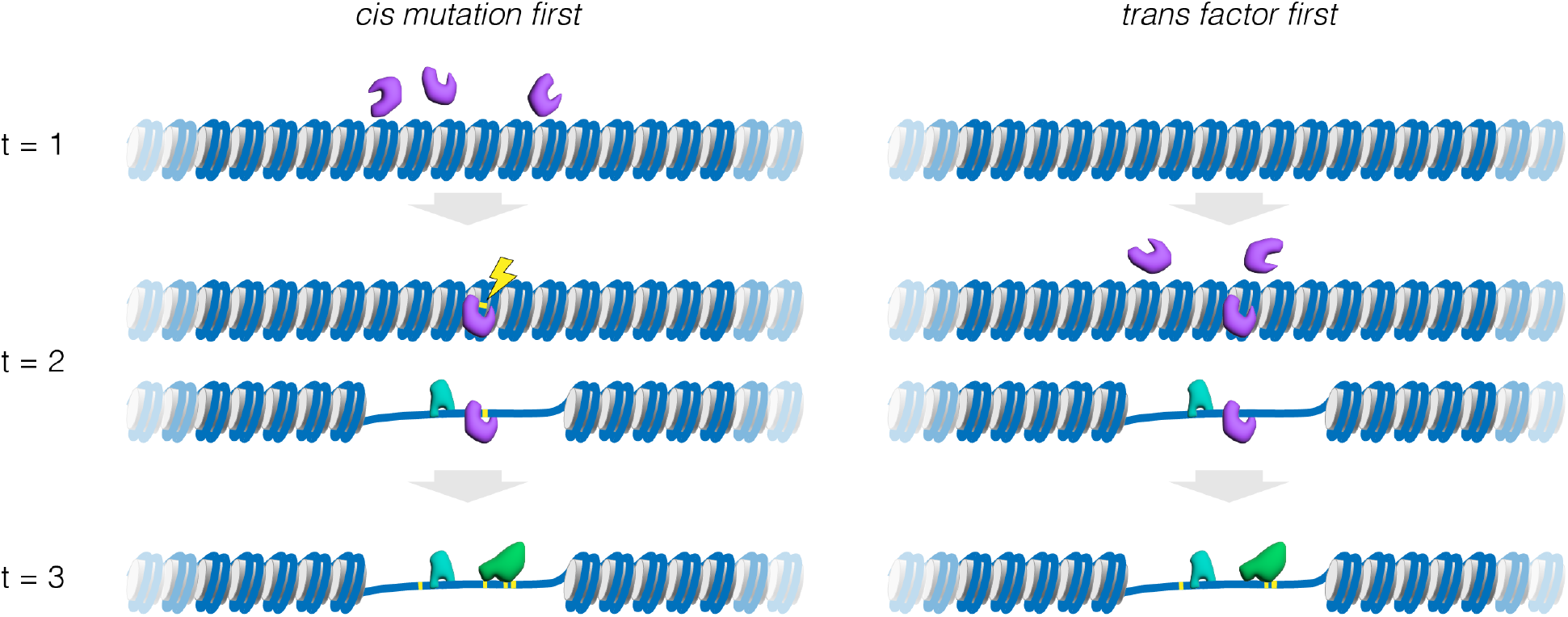
Mechanistic models for the evolutionary origin of an open chromatin region. Three distinct evolutionary time slices are illustrated: t=1 prior to the appearance of the open chromatin region (OCR), t=2 the event that precipitates local chromatin opening, and t=3 many generations after the origin of the OCR. Two general molecular processes could transform a given region of the genome from a relatively closed to a relatively open region of chromatin, based on the location of the genetic basis: *cis* or local and *trans* or distant. We illustrate this distinction based on a pioneer transcription factor, a class of proteins capable of sequence-specific binding to DNA in closed chromatin causing a local opening, typically on the scale of 1-2 nucleosomes (Zaret and Caroll, 2011). Note, however, that several other specific molecular mechanisms are also possible, involving chromatin remodeling complexes, histone acetylase/deacetylases, histone methylase/demethylases, and other proteins capable of acting to modify local chromatin configuration. (*A*) “*cis* mutation first” model. Here, the precipitating event is a local mutation (yellow; adjacent to the lightning bolt) that allows a pioneer transcription factor (purple protein) to bind to relatively closed chromatin. This pioneer factor was already present in the nucleus (see t=1), but can only bind locally following the new mutation that creates a binding site for it (t=2). Because pioneer factors can convert closed to open chromatin, other transcription factors also present in the nucleus (teal protein) may gain access to DNA within the newly opened region without the need for any additional mutations. (*B*) “*trans* factor first” model. Here, the pioneer factor is not originally present (t=1) in the nucleus at the developmental stage of interest. A mutation elsewhere in the genome (t=2) results in expression of the pioneer factor transcription factor, which can bind locally due to a pre-existing recognition site and thus without the requirement for a local mutation. Once present, the pioneer factor can open the local DNA, potentially allowing other transcription factors (teal protein) to bind without the need for any local mutations. Any impact on the transcription of a local gene could be susceptible to natural selection, since it has a genetic basis and a trait consequence. Over extended evolutionary timescales, additional mutations could accumulate within the OCR (t=3), potentially altering the suite of interacting transcription factors (for example, green protein now binds, purple protein no longer binds). In exceptional cases, natural selection may favor fixation of several new mutations within the OCR that alter gene expression. This figure illustrates the evolutionary origin of an OCR, but the evolutionary loss of an OCR could similarly occur through distinct molecular mechanisms based on the location of the genetic basis: for example, loss of a binding site for a transcription factor or loss of expression of a pioneer factor.

### Relationship of evolutionary regulatory mechanisms between proximal and distal elements

Next, we asked whether evolutionary changes in OCR sequence and chromatin accessibility differ between core promoters and more distal elements such as enhancers. We quantified changes in proximal (< 500 bp to translational start site of a gene) and distal (≥ 500 bp to translational start site of a gene) OCRs to test if either change covaries with promoter or enhancer evolution between species. While the transcription start site would be a more appropriate genomic reference for defining putative promoter regions, the reference genomes of these species currently lack annotation of 5’ untranslated regions for all genes. Thus, the translation start site was used instead to provide a consistent, albeit less sensitive, reference point of comparison of all genes for each species. Although many proximal and distal OCRs show evidence of positive selection on the *Ht* and *He* branches, we found no significant enrichment between the two categories of putative CREs (fig. 6b). This finding suggests mutations in both enhancers and promoters are likely important contributors in driving gene expression change, via altered binding and recruitment of regulatory factors and transcriptional machinery.

In contrast, accessibility changes show a much greater asymmetry between proximal and distal elements (fig. 6c). While accessibility distributions are comparable at early cleavage (16/32-cell embryos), many more accessibility differences accumulate in proximal OCRs relative to distal ones later in development, after zygotic activation and tissue specification has commenced. For example, by the gastrula and larval stages, more than 20% of proximal elements are significantly more accessible in *Lv* and *Ht*, whereas this figure is ∼5% for distal elements.

There are several possible biological and technical explanations for this result. Unlike enhancers, whose activity is often restricted to a single tissue or stage (Thurman, et al. 2012), promoters need to be accessible whenever the gene is transcribed, regardless of space and time. This has important implications for interpreting differences in chromatin configuration among species. At a biological level, greater promoter accessibility in the embryos of species with the ancestral life history (*Ht* and *Lv*) may merely reflect overall greater transcriptional activity and more changes in expression during development (fig. 1,3**; supplementary fig. 5)** (Israel, et al. 2016; Wang, et al. 2020). Secondly, for a given gene, the accessibility of a promoter region is normally greater than that of nearby enhancers (Thurman, et al. 2012), which may lend increased statistical power in detecting accessibility differences between species and contribute to an apparent enrichment of promoter accessibility changes. In addition, these chromatin configuration data were collected from whole embryos, so cell or lineage-specific differences in accessibility will be averaged within the sample, a technical factor that may be especially important for detecting accessibility changes of enhancer elements with tissue-specific dynamics. Lastly, promoter accessibility changes may simply be more often selected for when large changes in transcription are favored, at least for some genes. Adjusting enhancer accessibility can certainly induce changes in gene expression, but activating or shutting down a promoter may be a much more potent mechanism for large-scale changes in transcriptional regulation, such as those involved in rapid developmental evolution.

### Relationships between magnitude of gene expression and chromatin accessibility changes

Multiple lines of evidence presented above suggest that changes in the sequence or accessibility of *cis*-regulatory elements contributed to the evolution of development in *He*. The association between these changes and divergence in gene expression, however, is unclear. To understand this relationship, we compared mRNA expression fold-change of genes with accessibility fold-change of nearby OCRs at four developmental stages to measure instances of correlated changes in the magnitude of expression and chromatin configuration. A diagram of possible relationships between OCR accessibility and gene expression differences is provided in fig. 6d, to which datapoints are mapped and shown as density plots in fig. 6e. We partitioned the data into three categories: (1) all OCRs and genes (no filtering), (2) only differentially expressed genes and their nearby OCRs, and (3) differentially expressed genes and their nearby differentially accessible OCRs.

When all genes and OCRs are considered (fig. 6e: top row), there is no correlation between the magnitude and direction of accessibility and expression differences across developmental life histories. This result is not surprising at a global scale, given that expression levels and accessibility are conserved between species for most genes and OCRs, respectively. However, a notable set of proximal sites have greater accessibility in species with the ancestral life history at the blastula, gastrula, and larval stages, consistent with overall greater promoter accessibility in *Ht* and *Lv* (see previous section). A similar result occurs when only differentially expressed genes and nearby OCRs are considered (fig. 6e: middle row). One possible explanation for the lack of an obvious relationship is that, while only genes that are differentially expressed between life histories are plotted, each of these genes has an average of 3.1 OCRs assigned to it (based on proximity). Each of these OCRs may not be active at a given stage of development, or they may function antagonistically to one another (i.e., regulatory elements can enhance or repress transcription), thereby collectively dampening a correlative signal between gene expression and accessibility.

Lastly, we considered only differentially expressed genes and nearby OCRs with differential accessibility. This analysis reveals interesting relationships between transcription and chromatin configuration (fig. 6e: bottom row). As development progress, the relatively diffuse relationship between OCR accessibility and gene expression quickly tightens (compare 16/32-cell stage with larva). This likely reflects synchronization of zygotic genome activation with the embryo’s *cis*-regulatory landscape as maternal mRNA reserves are depleted and nascent transcription and cell differentiation commences. Greater accessibility at the majority of proximal sites in *Ht* and *Lv* is still found in this subset of genes and OCRs, though smaller sets of putative promoters with greater accessibility in *He* and corresponding expression increases are detected (faint gray spots). Among distal elements, we observed a considerably more diffuse relationship between accessibility, expression, and life history, possibly reflecting the potential activating or repressive functions of enhancers and downstream gene network interactions. At each developmental stage, the most common relationship was increased gene expression and accessibility in *Ht* and *Lv*, consistent with the preponderances of activating (as opposed to repressive) interactions within the ancestral dGRN.

Despite these relationships, our results highlight the difficulty in measuring correlations between the magnitude of cross-species changes in gene expression and regulatory element accessibility at a genomic scale. Prior studies have also reported a weak relationship between evolutionary changes in OCR accessibility and the expression of nearby genes between species (Shibata et al. 2012; Pizzollo et al. 2018). Several factors likely contribute a weak correlation, including: (1) a change in chromatin accessibility is just one of several possible molecular mechanisms that can change a gene’s expression; (2) individual CREs can have antagonistic effects on transcriptional regulation as activators or repressors, thus cancelling out effects; and (3) CRE accessibility is permissive rather than directive of regulatory factor binding and downstream gene expression, such that “openness” does not necessarily indicate the CRE is actively regulating transcription.

### Developmental trajectories of chromatin configuration and gene expression have evolved most frequently along the *H. erythrogramma* branch

To address the challenges associated with understanding the relationship between expression and accessibility changes of putative regulatory elements, we applied the “jump score” method to measure the magnitude of evolutionary changes in developmental trajectories, which are represented as distance in principal component space (Israel, et al. 2016). This metric is particularly useful because the jump score summarizes patterns of conservation or divergence in gene expression across development, rather than considering only a single stage. Here we apply this approach to evolutionary changes in the accessibility of putative CREs, as well as expression of genes, across developmental trajectories (fig. 8).

**Figure 8:**
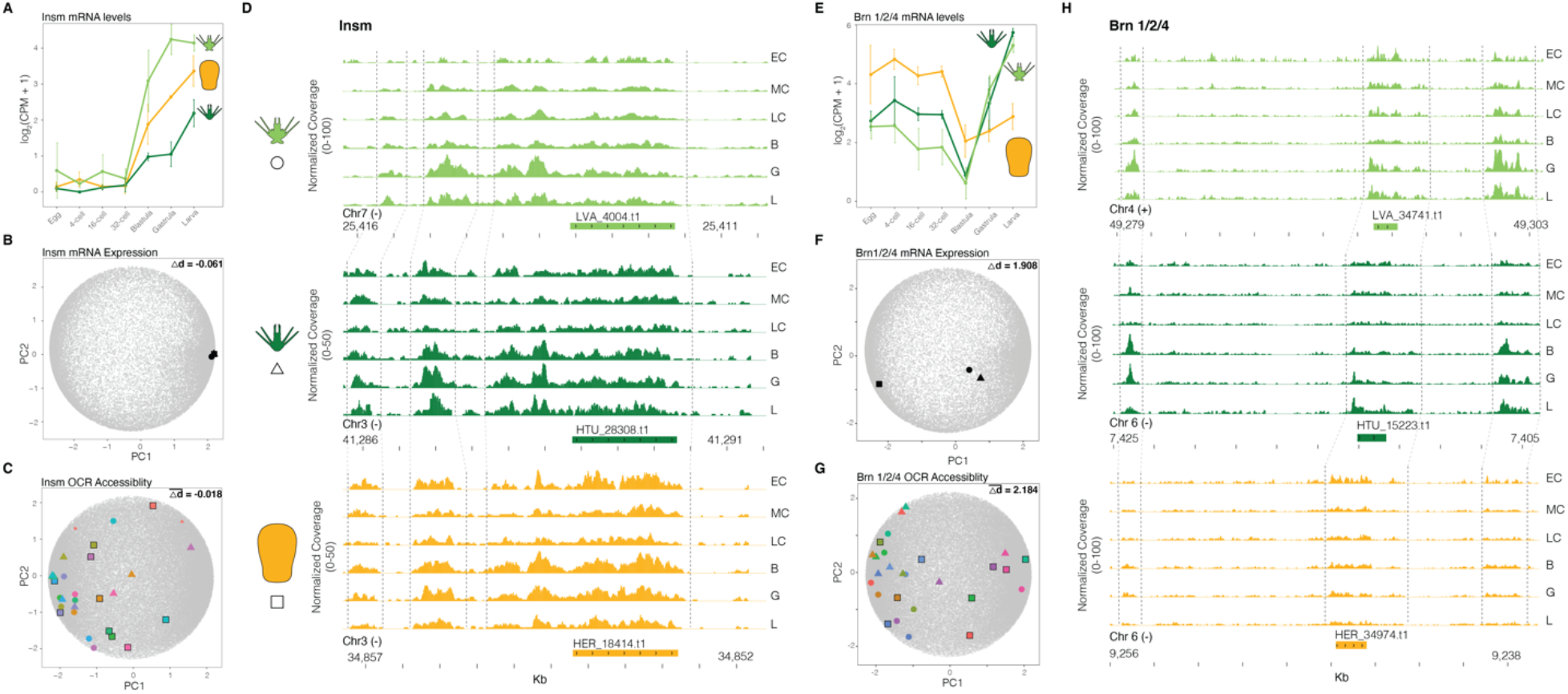
Quantifying developmental expression and accessibility changes in principal component space predicts divergent patterns of gene regulation. (*A-D*) An example of conserved developmental gene expression and chromatin accessibility for the gene *insm*. (*A*) Developmental mRNA expression profile of *insm* for each sea urchin species and (*B*) their coordinates in PCA space. (*C*) Coordinates in PCA space of developmental accessibility of orthologous OCRs near *insm* and (*D*) normalized chromatin accessibility coverage of these same OCRs through embryogenesis for each species. (*E-H*) An example of divergent developmental gene expression and chromatin accessibility associated with life history for the gene *brn 1/2/4*. (*E*) Developmental mRNA expression profile of *brn 1/2/4* for each sea urchin species and (*F*) their coordinates in PCA space. (*G*) Coordinates in PCA space of developmental accessibility of orthologous OCRs near *brn 1/2/4* and (*H*) normalized chromatin accessibility coverage of these same OCRs through embryogenesis for each species. See Supplementary Figure 6 for a summary figure of PCA distances for all genes and OCRs. OCR: open chromatin region; PCA: principal component analysis.

We compared the expression jump score of 9,423 genes and the mean jump score of nearby OCRs to test how developmental changes in transcriptional regulation and chromatin status are distributed along the *He* and *Ht* branches. For example, expression of *insm*, a transcription factor involved in neurogenesis in sea urchins (McClay, et al. 2018), is conserved across *He*, *Ht*, and *Lv* (fig. 8a). This is reflected by a low difference in jump scores between *He*-*Lv* and *Ht*-*Lv,* and tight clustering of expression profiles in principal component space (fig. 8b). Similar expression profiles are accompanied by conserved *cis*-regulatory landscape dynamics through development in each species, exemplified by a near-zero mean OCR jump score difference and accessibility profile of ten nearby OCRs (fig. 8c,d). In contrast, *brn 1/2/4*, a regulator of midgut patterning (Yuh, et al. 2005), has a divergent developmental gene expression trajectory in *He* relative to the ancestral state and large jump score differences in *He*-*Lv* and *Ht*-*Lv* branch comparisons (fig. 8e,f). This large expression change is accompanied by significant reduction in chromatin accessibility at the blastula through larval stages in *He*, represented by a large mean difference in OCR jump scores along the *He* branch (fig. 8g,h). This locus is especially interesting considering *He* larvae do not feed and have lost development of a functional through-gut.

Given the highly derived developmental program of *He* relative to the ancestral life history, we predicted that the most frequent changes in transcriptional regulation would occur along the *He* branch. Indeed, comparing jump scores of all genes and OCRs identifies concerted modifications to OCR accessibility and gene expression profiles during *He* development as the most frequent set of changes among genes and OCRs with divergent expression and chromatin accessibility trajectories (1,582 genes) (**supplementary fig. 6**; **Data S7**). Furthermore, the least common combination of changes includes both expression and accessibility changes occurring on the *Ht* branch (1,114 genes), consistent with an overall more conserved embryonic gene expression program in this species. Thus, we find that local patterns of CRE accessibility have evolved in concert with divergent developmental expression patterns during life history evolution in *He*. Collectively, our results suggest that modified transcription factor access to regulatory elements is associated with rapid and adaptive evolutionary changes to gene expression, and by extension, dGRN functionality. However, it remains to be determined which of changes evolved as a cause or consequence (or neither) of life history evolution in *He* — a challenge potentially addressable via empirical validation of candidate enhancer function.

### Distinct *cis*-regulatory modifications differentially effect evolution of gene expression

Lastly, we tested whether changes in nucleotide sequence and accessibility within OCRs are distributed near genes that are differentially expressed between developmental life histories. We found a small, though insignificant, increase in differential expression of genes with nearby differentially accessible OCRs relative to genes lacking differentially accessible OCRs at blastula, gastrula and larval stages (+6.2-9.9% change). No such increase exists during early cleavage (−1.8% change), which may be expected given the transcriptome is composed almost entirely of maternally-derived transcripts during early cleavage (Fisher’s exact test, two-sided: p > 0.05) (fig. 9a).

**Figure 9:**
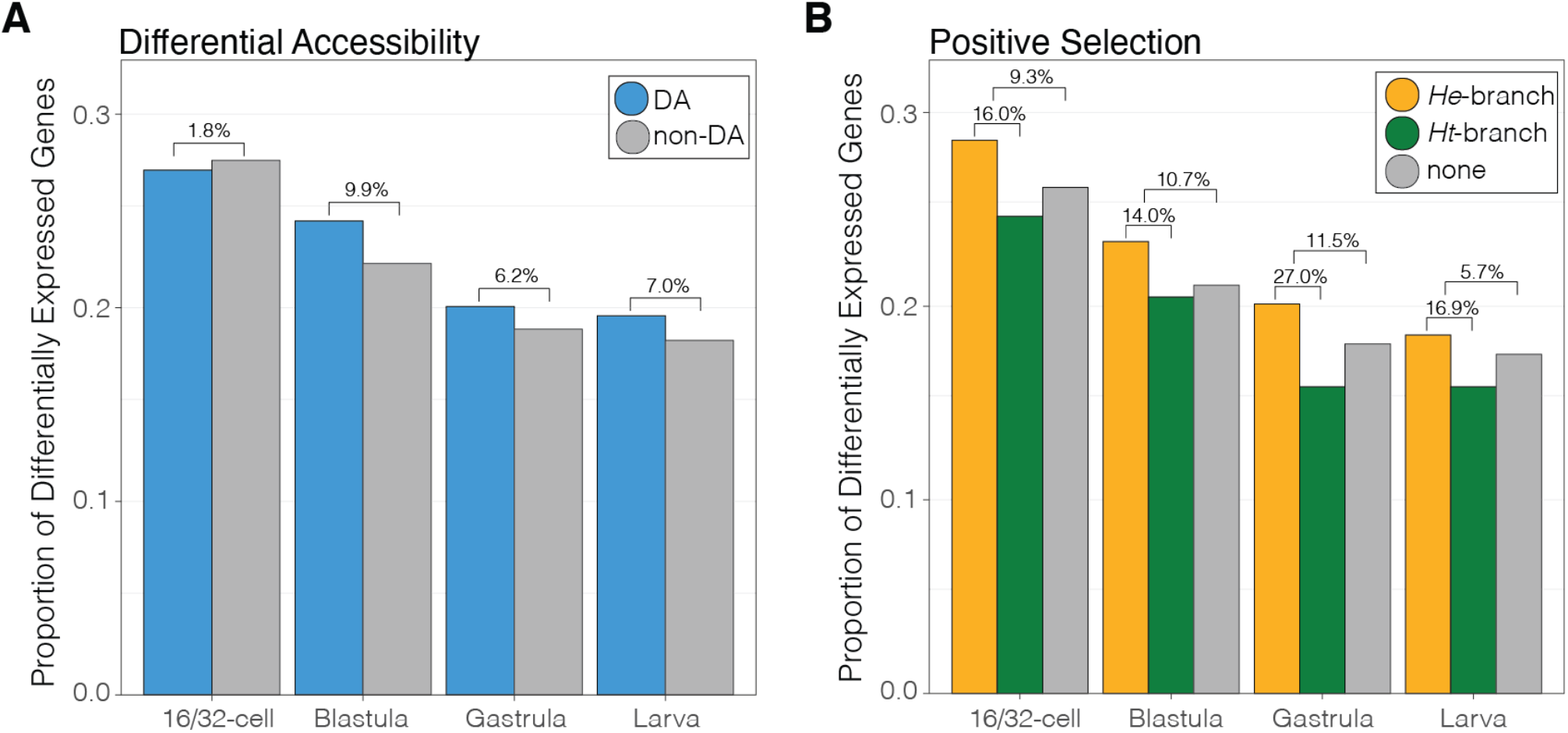
Evolutionary changes to *cis*-regulatory element function associated with life history evolution in sea urchins and expression changes of nearby genes. Proportion of genes that are differentially expressed between developmental life history strategies at four developmental stages, distinguished as having at least one nearby OCR with (*A*) differential accessibility or (*B*) evidence of lineage-specific positive selection. OCR: open chromatin region. Values above brackets indicate the percent change in proportion of genes that are differentially expressed between categories.

Similarly, we detected no significant difference in the proportion of differentially expressed genes and the presence of positive selection within OCRs of *He* when compared to OCRs lacking evidence of positive selection or OCRs with evidence of positive selection in *Ht* at each developmental stage (Fisher’s exact test, two-sided: p > 0.05) (fig. 9b). Still, we note that throughout development, incidences of differential expression are higher when there exists one or more nearby OCRs with evidence of positive selection relative to OCRs lacking evidence of positive selection (+5.7-11.5% change; fig. 9b: orange and gray bars) or OCRs with evidence of positive selection in *Ht* (+14.0-27.0% change; fig. 9b: orange and green bars). This finding is concordant with our predictions of evolutionary mechanisms of gene expression change, as selective changes in CRE sequences in *He* are most often coupled with concordant divergence in developmental gene expression possibly associated with their derived life history strategy. Positive selection within CRE sequences of *Ht*, on the other hand, are less often linked to changes in gene expression associated with life history, as *Ht* has retained the ancestral sea urchin developmental program. Lastly, while the absolute number of accessibility changes between developmental life histories is typically greater than detected instances of positive selection (figs. 3,5), mutations within putative CRE sequences are more often associated with changes in gene regulation and expression relative to a no-change state (mean +9.3% change) than changes in accessibility (mean +5.3% change).

These results may reflect distinct characteristics of the molecular mechanisms that modify *cis*-regulatory function. In particular, an evolutionary change in the chromatin accessibility of CRE might only manifest at certain developmental stages or in certain cell types, whereas a change in the DNA sequence of a CRE is present throughout development and in every cell type. For this reason, mutations that influence chromatin status might in general have lower pleiotropy than mutations within OCRs. This possibility is an extension of the idea that mutations in CREs that drive tissue-specific expression may have lower pleiotropy than those in CREs that influence expression throughout development and in all cell types (Wray et al. 2003; Stern 2010; Singh and Yi 2021; but see Preger Ben-Noon et al. 2018). Evolutionary changes in chromatin configuration are an additional molecular mechanism with the potential to limit pleiotropy, by physically masking a mutation within a CRE such that it can only exert an effect on transcription when chromatin is open.

Conversely, opening chromatin in a larger range of conditions could expose cryptic genetic variation to natural selection. Although there is good empirical evidence that cryptic genetic variation exists within natural populations (Gibson & Dworkin, 2004; Paaby and Rockman, 2014), it is less clear how it becomes visible. One possibility (among others) is that mutations accumulate within a cell type-specific CRE that have no consequence because they do not influence interactions with any of the *trans*-acting factors in that particular cell type. An evolutionary change that opens the local chromatin in a different cell type might, however, expose those new mutations to a different suite of trans-acting factors, potentially altering their interactions with the CRE and thus transcription. Deconvolving the interplay and evolutionary consequences of epigenetic and sequence changes within CREs remains an important challenge for understanding the evolution of transcriptional regulation.

## Conclusions

In this study, we leveraged a recent life history switch in sea urchins that was accompanied by elevated rates of adaptive changes in developmental gene expression (Israel et al. 2016; Wang et al. 2020; Wray 2022) to investigate how evolutionary mechanisms act on mechanisms of transcriptional regulation and embryonic gene expression. In particular, we distinguish two distinct molecular mechanisms within *cis*-regulatory elements that could alter gene expression: sequence changes that might influence transcription factor:DNA interactions and epigenetic changes that might limit access of transcription factors to DNA. We find that evolutionary changes in chromatin are common near differentially expressed genes, especially on the branch leading to the derived life history and especially so near critical developmental gene regulatory network (dGRN) genes (fig. 10, left column). Elevated accumulation of mutations within OCRs indicative of positive selection is not enriched on the branch leading to the derived life history when considering all genes, but strikingly enriched near dGRN genes specifically (fig. 10, central column). These distinct molecular mechanisms may also act synergistically, as seen by enrichment for both changes within putative dGRN CREs on the branch leading to the derived life history (fig. 10, right column). Although extensive prior evidence indicates that changes in protein:DNA binding contributes to the evolution of gene expression, much less empirical support exists regarding the contribution of epigenetic changes. Our results suggest that evolutionary changes in local chromatin configuration during early development can contribute to the evolution of embryonic gene expression, while branch- and dGRN-specific enrichments suggest that many of these changes are adaptive and may be tied to the evolution of the derived life history. More broadly, the comparative approaches developed in this study offer new ways to analyze multimodal functional genomic datasets to investigate the evolutionary mechanisms that operate on *cis*-regulatory elements and their consequences for gene expression and organismal traits.

**Figure 10:**
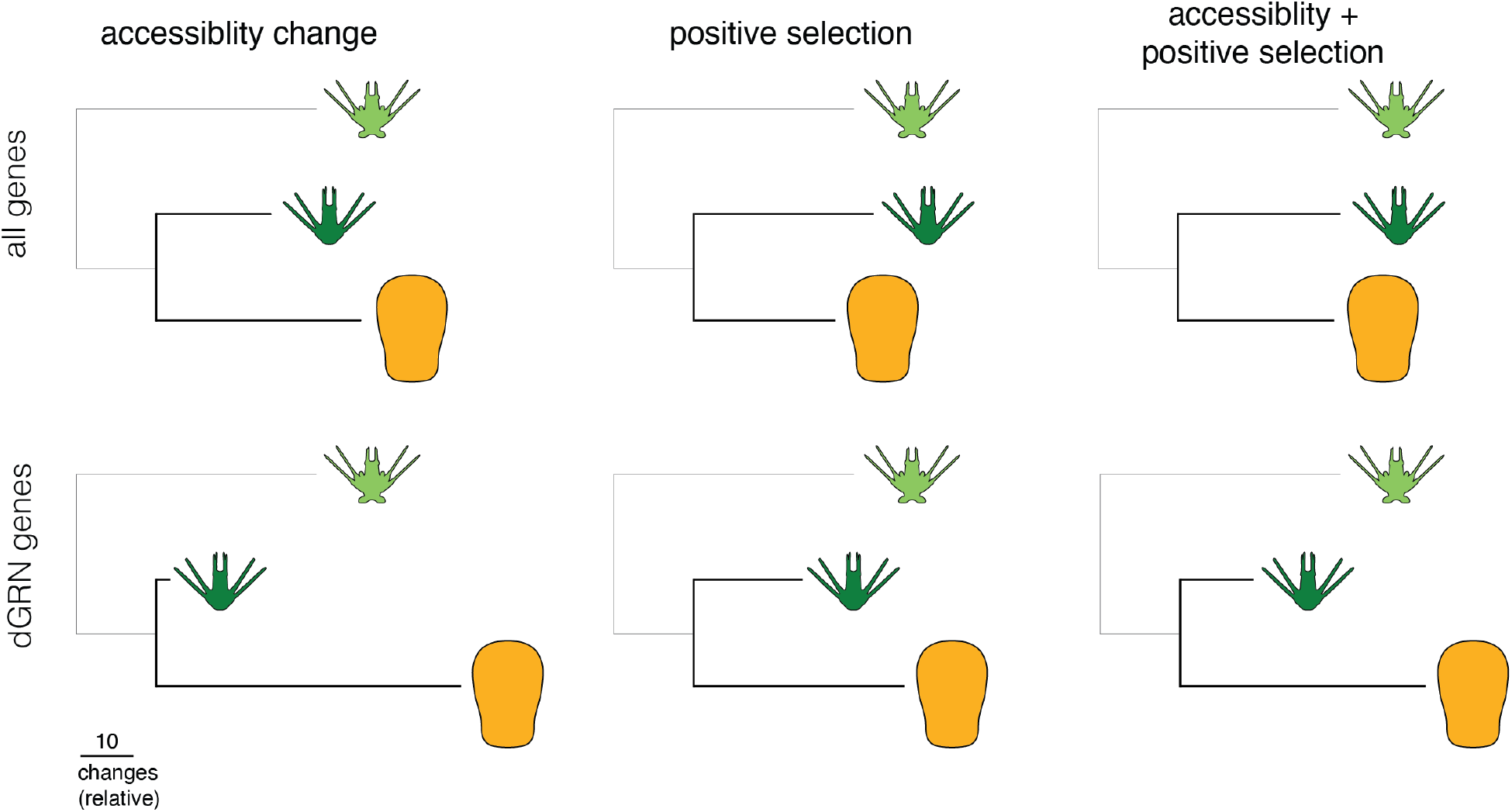
Chromatin accessibility changes, mutational signatures of positive selection, and the coincidence of both evolutionary changes are abundant in the *H. erythrogramma* genome, especially nearby dGRN genes. Enrichment of regulatory modifications to OCRs near dGRN genes suggests selective pressures imposed by changes in developmental life history can rapidly modify *cis*-regulatory interactions underlying developmental processes. Tandem accessibility and mutational changes may act as a common gene regulatory method to efficiently modify gene expression for organismal adaptation and innovation. Branch lengths scaled to relative number of changes between *Heliocidaris* branches for each set of modifications.

## MATERIALS AND METHODS

### Embryo Rearing and Sample Preparation

*H. erythrogramma* and *H. tuberculata* adult animals were collected near Sydney, NSW, Australia and maintained at 21°C in aquaria with circulating natural seawater. *L. variegatus* individuals were collected near the Duke University Marine Lab in Beaufort, NC, USA and maintained at 23°C in aquaria at Duke University, Durham, NC, USA. Each sea urchin species was induced to spawn via injection of 0.5 M KCl solution into the coelom. Eggs were washed twice in 20 µm filtered seawater (FSW). Sperm was collected dry and stored at 4°C until usage. Washed eggs were fertilized with 1 µL of sperm, and after 10 minutes, were washed three times in FSW. For each species, three unique male-female pairs were crossed to produce three biologically-independent replicates of sea urchin embryos. Embryos were reared at 23°C in large glass culture dishes supplied with FSW, and water was changed every six hours. Because these species exhibit different developmental rates, developmental milestones, rather than absolute developmental time, were selected to maximize developmental synchrony within cultures and between species for comparison. These stages include 1) 4^th^/5^th^ (EC), 2) 6^th^/7^th^ (MC), and 3) 8^th^/9^th^ (LC) cleavage stage embryos, as well as 4) hatched blastula (B), 5) mid-gastrula (G), and 6) early larva (L) (two-armed pluteus for *H.* tuberculata and *L. variegatus*; 32 hours post-fertilization for *H. erythrogramma*). Quickly distinguishing embryonic stages is difficult for cleavage stage embryos, so we closely monitored development of each culture and used known division rates for each species and water temperature to estimate when the culture would be most enriched for the targeted developmental stage. Once a culture reached the appropriate stage, a subset of the culture was retrieved for processing, stage confirmed by visual inspection, arrested or delayed embryos removed by hand, and the remaining embryos transferred into a 1.5 mL Eppendorf tube for ATAC-seq processing. Microscopic examination confirmed that most embryos had reached the target stage.

### ATAC-seq Protocol and Sequencing

ATAC sample preparation was carried out according to the Omni-ATAC-seq protocol (Corces, et al. 2017). For each replicate, embryos were washed once in 1 µm filtered seawater, lysed, and ∼50,000 nuclei were isolated for the transposition reaction using the Illumina TDE1 enzyme and tagmentation (TD) buffer (Cat. No. 20034197 and 20034198) (San Diego, CA, USA). Sequencing libraries for each replicate were amplified via PCR after separately determining the optimal number of amplification cycles for each sample using qPCR as described in (Buenrostro, et al. 2015), and libraries were purified and size selected using Ampure XP Beads at a 1.8:1 bead volume:library volume (Beckman Coulter, Brea, CA, USA). Library quality and transposition efficiency was accessed using a Fragment Analyzer and PROSize 2.0 (Agilent, Santa Clara, CA) at the Duke University Center for Genomic and Computational Biology. *H. erythrogramma* and *L. variegatus* libraries were sequenced on an Illumina HiSeq 4000 instrument using 50 bp SE sequencing. *H. tuberculata* libraries were sequenced on an Illumina NovaSeq 6000 instrument using 50 bp PE sequencing (only SE were used for data analysis).

### ATAC-seq Data Analysis

Raw ATAC-seq reads were trimmed for quality and sequencing adapters using “cutadapt” (Martin 2011) v2.3 with the following parameters: -a CTGTCTCTTATACACATCT -q 20 --trim-n -m 40. Trimmed reads were then aligned to each species’ respective genome via “stampy” (Lunter and Goodson 2011) v.1.0.28 using the “—sensitive” set of parameters. The *Lv* genome was retrieved from Davidson et al. 2020 (NCBI BioProject: PRJNA657258). The *Heliocidaris* genome assembles were retrieved from Davidson et al. 2021 (NCBI BioProject: *H*e-PRJNA827916; *Ht*-PRJNA827769). ATAC-seq alignments were filtered for mitochondrial sequences and required an alignment quality score of at least 5 using “samtools” (Li, et al. 2009) v.1.9. Filtered alignments are available in BAM and Bedgraph format (see “Data Availability” below).

In this study, we aimed to compare evolution of orthologous regulatory elements so to accomplish this, we performed a series of liftOvers (Hinrichs, et al. 2006) to convert ATAC-seq alignments between genomic coordinates of each sea urchin species. We took an iterative, reciprocal strategy described below to minimize possible reference bias associated with converting between genome assemblies, requiring a minMatch of 0.75 when lifting alignments between species pairs: 1) *H. erythrogramma*: *He* → *Lv* → *He*; 2) *H. tuberculata*: *Ht* → *Lv* → *He*; 3) *L. variegatus*: *Lv → He* → *Lv* → *He*. At the end of filtering and coordinate conversion, all ATAC-seq alignments were referenced to the *H. erythrogramma* genome. LiftOver files are available in (Davidson, et al. 2021).

ATAC-seq peaks were called from these filtered, converted alignments using the “MACS2” v2.1.2 (Zhang, et al. 2008) callpeak function (parameters: –nomodel, –keep-dup=auto, –shift 100, –extsize 200) for each developmental stage separately at an FDR threshold of 5%. Peak coordinates were merged using the “bedtools” (Quinlan and Hall 2010) v2.25 *merge* function requiring a peak overlap of at least 200 bp to be merged into a single peak. Lastly, for each sample, accessibility of each peak was measured with the “bedtools” *multiBamCov* function and imported into R v4.0.2 for statistical analysis (Data S1). While not a primary focus of this study, all ATAC-seq peaks and raw accessibility counts for each species, independent of sequence conservation between species, is additionally provided in Data S8, as are ATAC-seq peaks specific to *Lv* (not found in *Heliocidaris*).

### Estimates of positive selection

To measure branch-specific signatures of positive selection in the non-coding genome, the “adaptiPhy” (Berrio, et al. 2020) pipeline (https://github.com/wodanaz/adaptiPhy) for global tests of natural selection was followed. First, orthologous sequences for non-coding sites of interest (ATAC-seq peaks) were extracted from each species’ genome into FASTA format. Sequences were trimmed to include only contiguous DNA sequence using the *prunning.py* script and filtered using the *filtering.py* script, requiring a minimum alignment length of 75 base pairs. These trimmed and filtered alignments serve as “query” sequences of tests for selection. To generate a neutral reference for comparison, ten orthologous, neutral sites were randomly selected from a curated set of ∼88,000 putatively neutral sequences (Davidson, et al. 2021) (Data S2) and concatenated into a single neutral reference sequence. A global neutral reference, rather than a local reference, was implemented to more conservatively estimate signatures of positive selection as recent work has demonstrated a local neutral reference may more likely overestimate positive selection – an especially acute issue in smaller, gene-dense genomes (Berrio et al., 2020). Then, substitution rates of both the query and randomly concatenated neutral reference were estimated using phyloFit (Siepel and Haussler 2004), and the zeta score was calculated as the ratio of the query substitution rate to the neutral reference substitution rate. In addition, p-values of likelihood ratio tests for significant levels of branch-specific positive selection were calculated with adaptiPhy (Berrio, et al. 2020) pipeline using HyPhy (Pond, et al. 2005). For each query site, tests for positive selection were repeated 10 times against a unique neutral reference to ensure reproducible estimates of zeta and significance values for each non-coding site of interest. P-values and substitution rates for all query and neutral sites were then imported in R v4.0.2. For each query site, the median zeta score and p-value for each test of positive selection across the ten replicate calculations were used for downstream analyses.

### OCR Filtering

After accessibility and estimates of positive selection were calculated for each ATAC-seq peak, herein referred to as an “open chromatin region” (OCR), a series of filtering and quality control metrics were carried out to ensure only high confidence and quality peaks were compared between species. These filtering steps are as follows: 1) each OCR is required to have at least 75bp of contiguous, single copy sequence (see previous section) for accurate estimations of selection ; 2) a local composition complexity (LCC) (Konopka 2001) value of 1.9 or more was required of the OCR for each species to remove repetitive or other low-complexity sequences that may generate inaccurate estimations of selection (module: biopython.org/docs/1.75/api/Bio.SeqUtils.lcc.html); 3) a CPM of 3 or more was required in at least 9 samples to remove OCRs with extremely low accessibility across samples; 4) the midpoint of the OCR must lie within 25 Kb (in either direction) of the translational start site of a gene model. The 25 Kb window for assigning OCRs to genes was selected on the basis that the median and mean intergenic distance within the *He* genome are 21.4 Kb and 31.9 Kb, respectively. See Supplementary Table 1 for a summary of these filtering steps and resulting number of OCRs. Given nearly no prior knowledge is known of the *cis*-regulatory landscape for these sea urchin species, these stringent filtering methods were carried out in order to maximize confidence in comparisons of non-coding sequence evolution and function. This resulted in a final set 35,788 high-confidence OCRs for cross-species analysis (Data S3).

### ATAC-seq Statistical Analysis

Principal component analysis and variance partitioning was performed on the regularized log-transformed (rlog) data (Love, et al. 2014) of filtered OCR accessibility counts via the *prcomp* function and the “variancePartition” v1.18.3 package (Hoffman and Schadt 2016) in R using the formula: ∼ (1 | Stage) + (1 | Species) + (1 | Life History) + (1 | Genus) + (1 | Stage: Life History). Raw accessibility counts of the filtered OCRs were loaded into DESeq2 (Love, et al. 2014) v1.30 to calculate differential accessibility between sample groups. Differentially accessible sites between life history strategies were classified as having a 2-fold accessibility difference and supported by an FDR of 10% in *He* vs *Ht* and well as *He* vs *Lv* comparisons (including the sign of the accessibility difference being in the same direction) (see **supplementary fig. 7** for all significance cutoff combinations of differential accessibility results). For graphical purposes, the average fold-change in the *He* vs *Ht* and *He* vs *Lv* comparisons is displayed within figures. Significant levels of positive selection in either the *He* or *Ht* branch were classified as OCRs having a zeta value greater than 1.25 and supported by a median false-discovery rate less than 10% in one species, and failing to meet both criteria in the other. Motif enrichment analyses were carried out with “Homer” v4.11 (Heinz, et al. 2010) via the findMotifsGenome.pl script using all OCRs as the background set and regions size of 200 bp. Sea urchin GRN genes are provided in Data S4, retrieved from the Biotapestry project of the Institute of Systems Biology, Seattle, WA, USA (www.biotapestry.org: accessed June 27, 2017).

### RNA-seq Expression Analyses

Raw RNA-seq reads from eggs, 4-cell, 16-cell + 32-cell (collectively EE), blastula (B), gastrula (G), and early larva (L) stage embryos of *He*, *Ht*, and *Lv* were retrieved from (Israel, et al. 2016), trimmed and filtered for low quality bases and reads with Trimmomatic (Bolger, et al. 2014), and aligned to each species respective genomes and gene models with STAR (Dobin, et al. 2013). From these alignments, mRNA expression was estimated with Salmon (Patro, et al. 2017) and loaded to R for statistical analysis. Read counts for summed to each gene’s best match to the *S. purpuratus* v4.2 gene models to generate a common reference for expression comparisons between species as described in (Israel, et al. 2016) (Data S5). Differentially expressed genes between life histories were called as having a fold-change in expression > 2 and supported by an FDR of 10% or less in *He* vs *Ht* and well as *He* vs *Lv* comparisons (including the sign of the accessibility difference being in the same direction) (Data S6). These criteria were chosen to make RNA and ATAC-seq analyses between species comparable as possible.

### Principal Component Distances of Developmental Profile

To identify OCRs and genes whose developmental regulatory profiles diverged in *H. erythrogramma* or *H. tuberculata*, we calculated principal component distances of chromatin accessibility and gene expression profile shapes between species throughout development as described in (Israel, et al. 2016). In this method, developmental time course data is plotted onto principal component space for each species, and pairwise distances between species are calculated for each gene or OCR (see below). Large differences in these distances provide evidence that the shape of the developmental trajectory for a given gene or OCR has changed in one of the species (the species with the larger pairwise distance). Mean accessibility and expression profiles of regularized log-transformed (rlog) OCR and genes counts were standardized using the *standardize* function of the “mfuzz” v2.48.0 package (Kumar and M 2007). From these data, scaled distance measures (“jump scores”) between orthologous non-coding sites or genes were calculated using loadings of the first two principal components (Data S7): 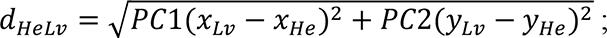 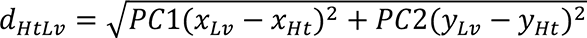. A significant change in the developmental trajectory of gene expression or OCR accessibility was determined to have a jump score difference of at 0.15 on one branch over the other.

## DATA AVAILIBILITY

Raw ATAC-seq reads are available in FASTQ format on NCBI’s Short Read Archive (PRJNA828607). Filtered alignments of ATAC-seq reads before and after between-species liftOver files are available on Dryad (https://doi.org/10.5061/dryad.jsxksn0cp). Results files of ATAC-seq analyses are available as Supplementary Data files.

## AUTHOR CONTRIBUTIONS

PLD and GAW conceived and designed the study. PLD and MB performed sample collection. PLD performed the data analysis. PLD and GAW wrote the manuscript. All authors revised the manuscript.

## ACKNOWLEDGEMENTS

We would like to thank the Sydney Institute of Marine Sciences and their staff for assistance with animal care, laboratory set up, and carrying out experiments. We also thank Allison Edgar and Hannah Devens for assistance preparing sea urchin embryo cultures associated with this work. This work was supported by the National Science Foundation Division of Integrative Organismal Systems (award no. 1929934 to GAW).

## COMPETING INTERESTS

The authors have no competing interests to declare.

## Notes

https://doi.org/10.5061/dryad.jsxksn0cp

### Competing Interest Statement

The authors have declared no competing interest.

### Summary of Updates

Improved interpretation of ATAC-seq results, revised Figure 3, and more thorough explanation of models and conclusions

