## Supplementary Fig. for "Evolutionary changes in the chromatin landscape contribute to reorganization of a developmental gene network during rapid life history evolution in sea urchins"

for

| Filter | *Description* | *# OCRs* |
| --- | --- | --- |
| None | Initial number of OCRs | 254,644 |
| CPM | At least 9 samples (equivalent to 3 sample groups) required to have a CPM ≥ 3 | 135,754 |
| Orthology | OCR’s closest gene must match in *Lv*, *Ht*, and *He* and must be 1-1-1 between species | 49,545 |
| Adaptiphy | Orthologous OCRs must have ≥ 75 bp of contiguous DNA in each species | 45,651 |
| Low complexity | OCRs must have a local complexity composition score ≥ 1.9 in each species | 39,982 |
| OCR-Gene Distance | Midpoint of OCR must lie within 25 kbp of translation start site of a gene | 35,788 |

**Supplementary Table 1:** Results from filtering steps of orthologous OCRs identified in this study, including the type of filter, the filter description, and the number of OCRs remaining after implementing the filter.

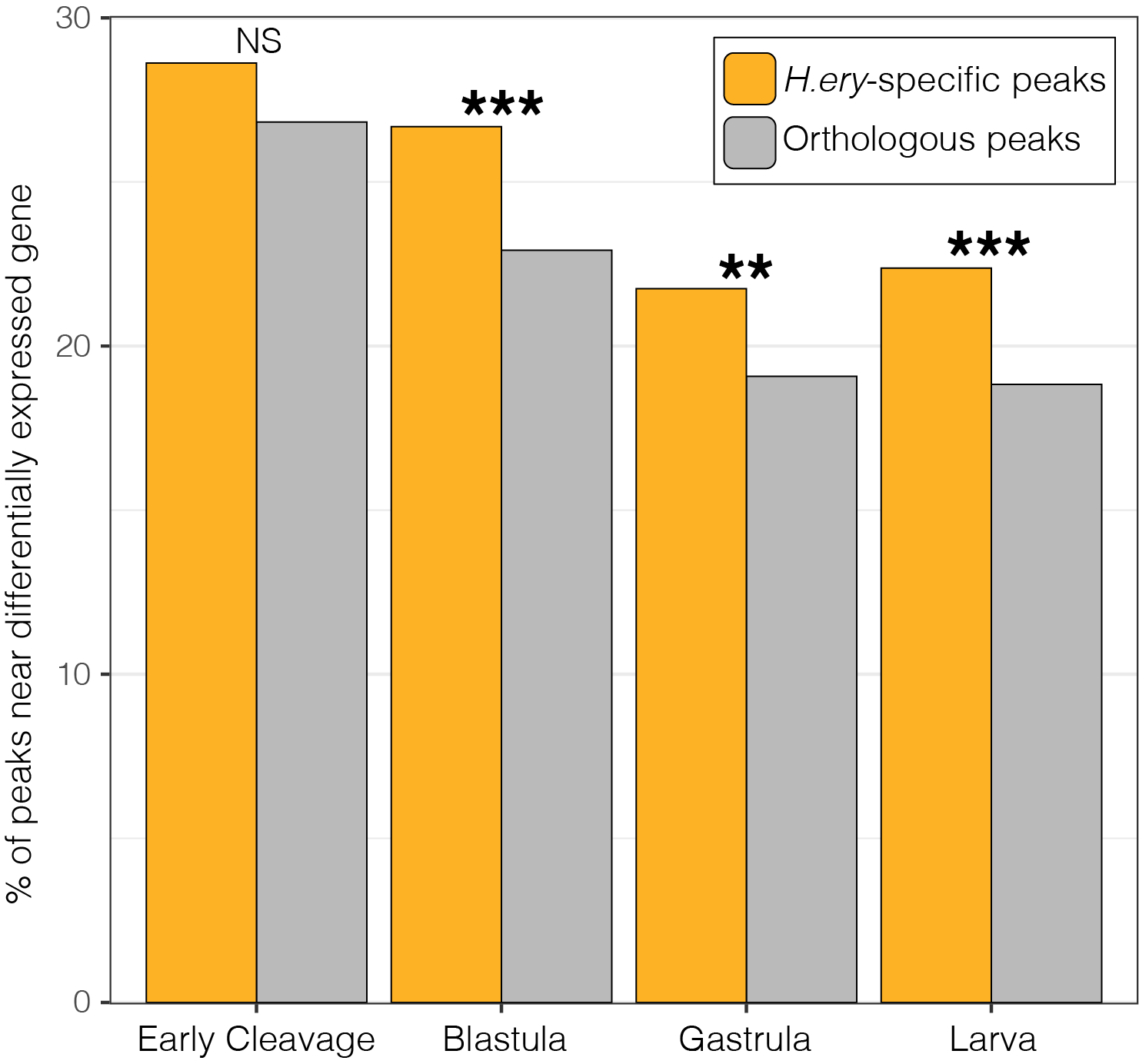

**Supplemental Figure 1:** Proportion of OCRs near a differentially expressed gene that are only found in the *H. erythrogramma* genome (“*H.ery-*specific”) or found in all three species analyzed in this study (*Lvar*, *Htub*, and *Hery*) (“orthologous”). Proportions of *H.ery*-specific OCRs tends to be greater than proportion of orthologous OCRs near differentially expressed genes, significantly so at the blastula through larval stages. ** and *** indicates p-value ≤ 5e-3 and p-value ≤ 5e-4 in Chi-square test of independence, respectively.

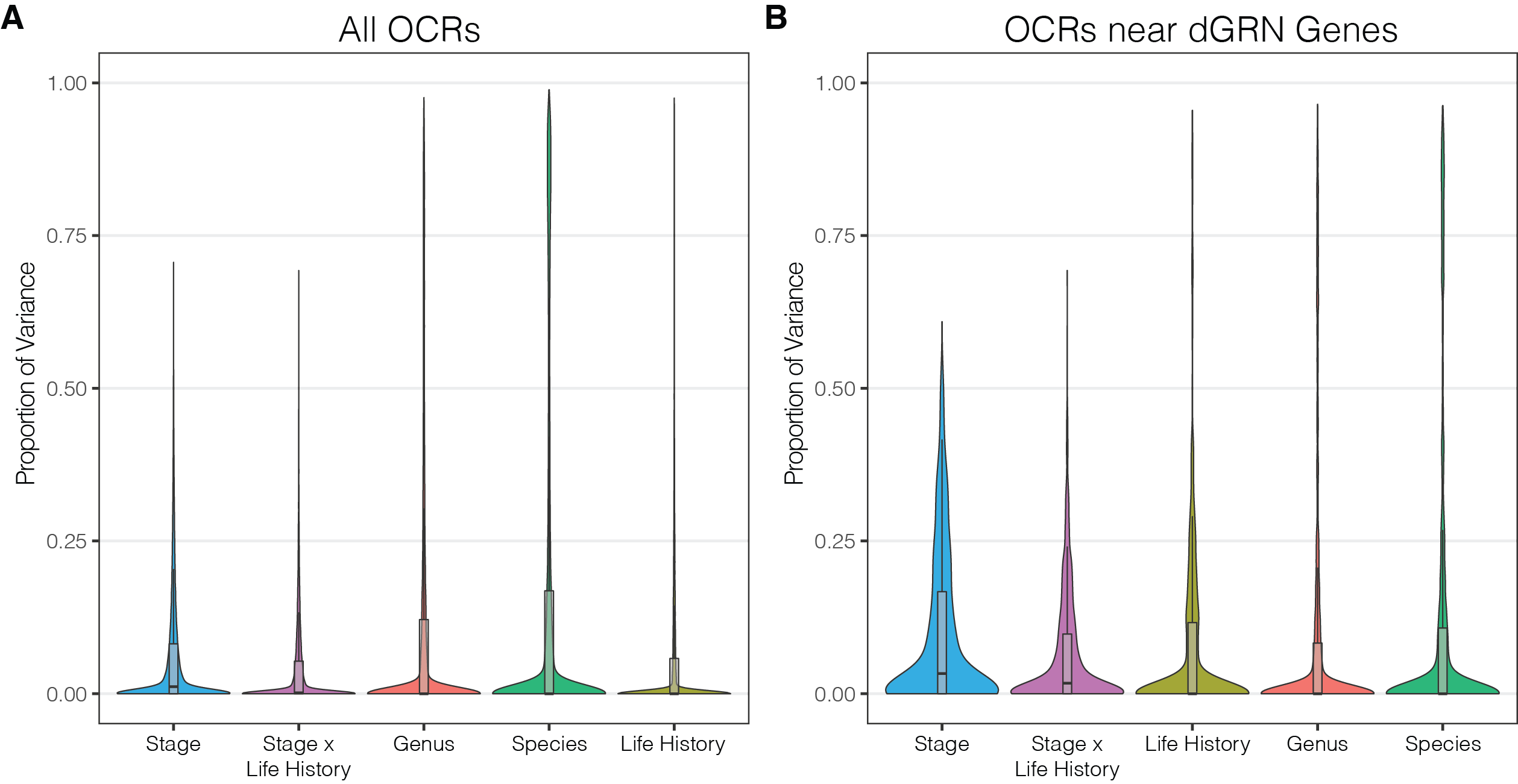

**Supplemental Figure 2:** Results of variance partition analysis of open chromatin region (OCR) accessibility. Differences in developmental life history strategy contributes disproportionally to chromatin accessibility near developmental gene regulatory network (dGRN) genes relative to the entire genome.

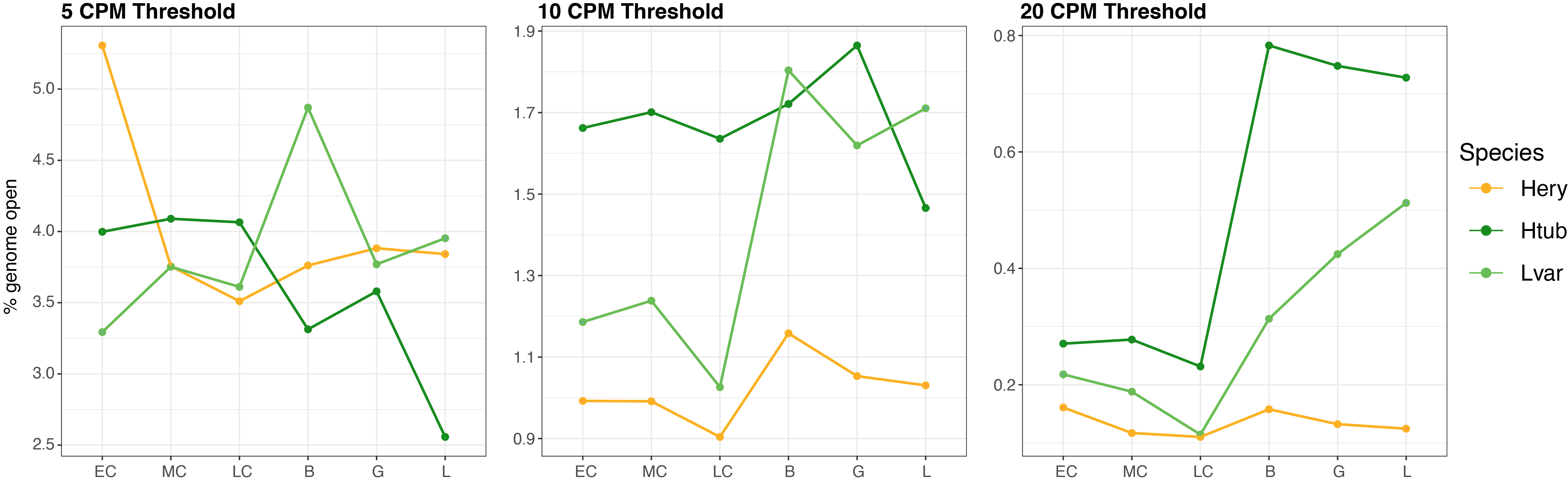

**Supplemental Figure 3:** Percent of each species’ genome that is “open”, determined by significantly open chromatin region peaks from ATAC-seq data. CPM threshold indicates ATAC-seq count required in order to be considered a significantly “open” peak. This requirement had to be met in each sample from each developmental stage analyzed. Approximately equal proportions of “open” base pairs between species at a 5 CPM threshold likely reflects inclusion of small, more variable peaks and thus may be less likely accurately capturing biological signal in this calculation.

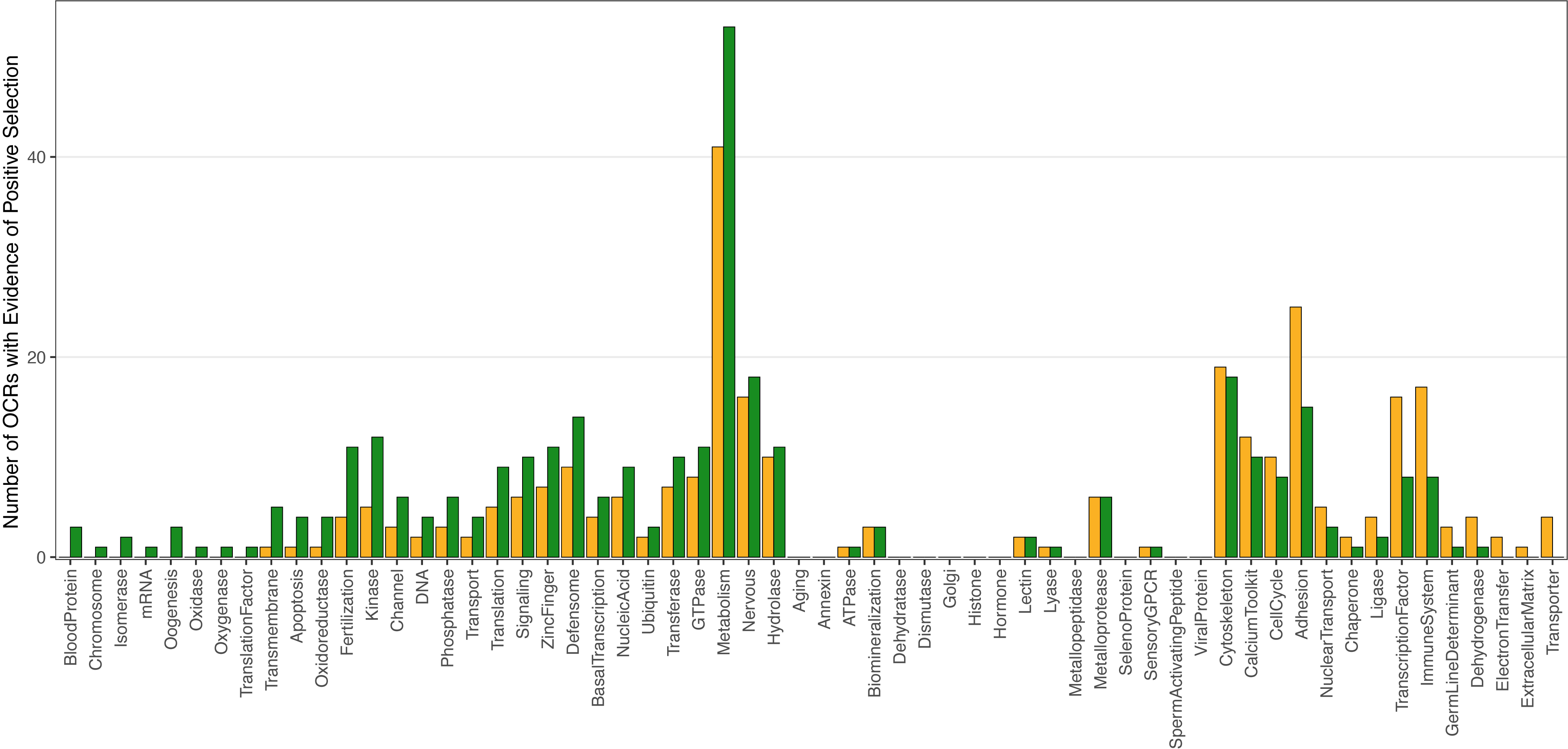

**Supplemental Figure 4:** Number of OCRs with evidence of positive selection on either the *H. erythrogramma* (orange) or *H. tuberculata* (green) branch, partitioned by proximity genes of various functional categories.

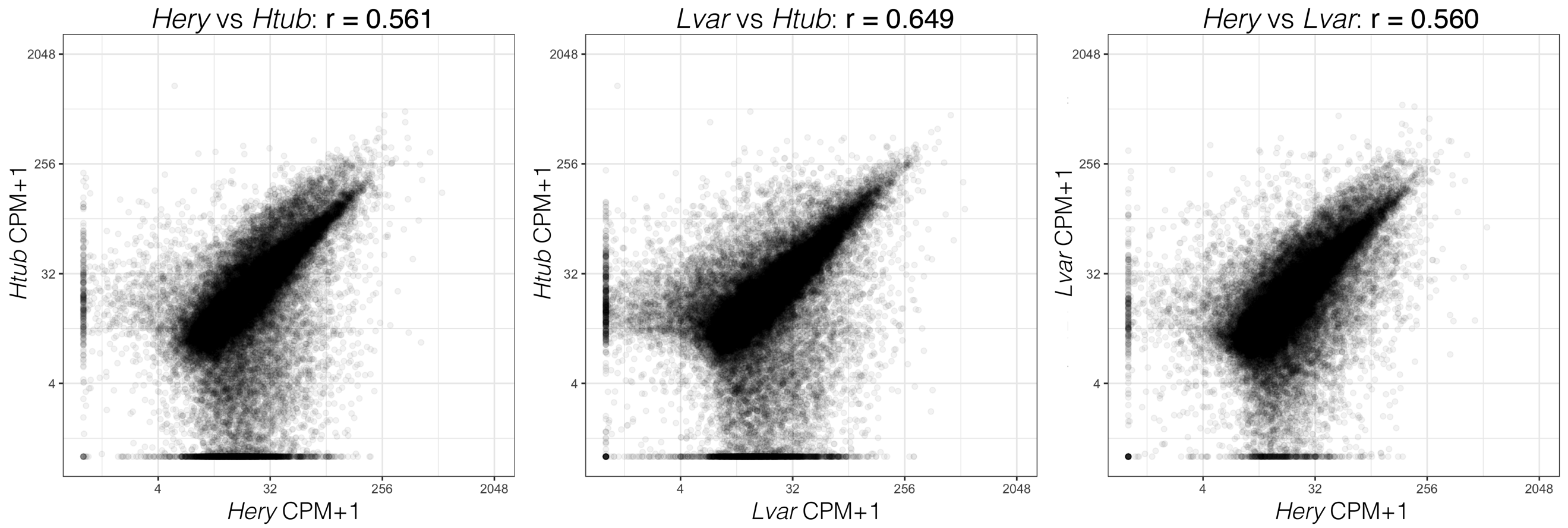

**Supplemental Figure 5:** Correlation of OCR accessibility between each species pair, summarized across developmental stages. Higher overall correlation is measured between planktotroph accessibility (*Htub* and *Lvar*) despite the much longer divergence time between these two species (~ 40 my) relative to the *Heliocidaris* species (~ 5 my).

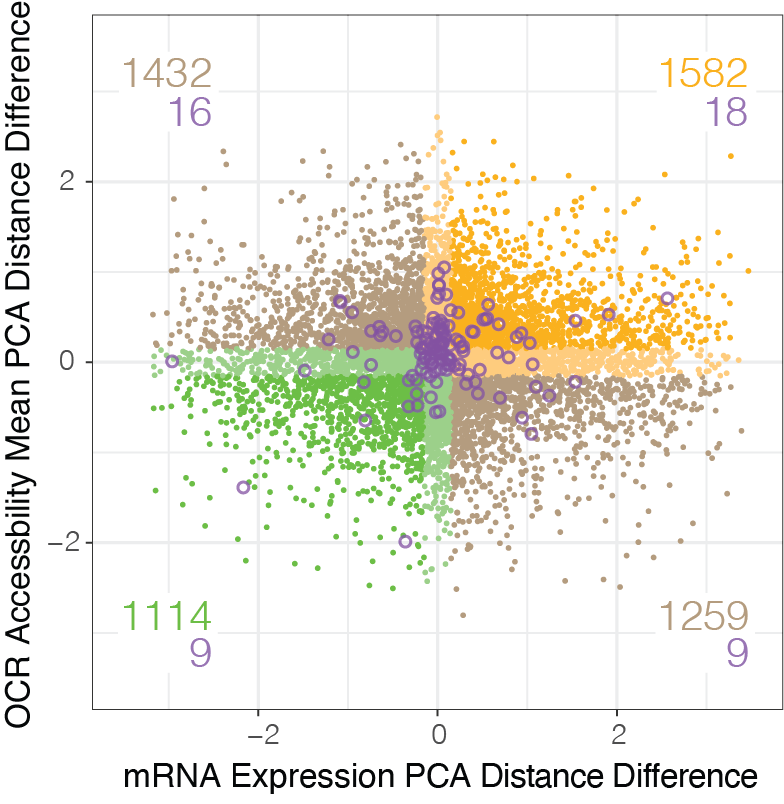

**Supplemental Figure 6:** Principal component analysis (PCA) distance differences of gene expression and mean chromatin accessibility developmental profiles. Distance differences are calculated as the difference between *He* and *Lv* vs *Ht* and *Lv* in PC space (see *Methods* and Data S7). Developmental gene regulatory network genes denoted by purple circles. Orange points indicate genes whose chromatin accessibility and gene expression profiles diverge concordantly along the *He* branch, green points indicate genes whose chromatin accessibility and gene expression profiles diverge concordantly along the *Ht* branch, and brown points indicate genes that have discordant divergence in accessibility and gene expression between either branch.

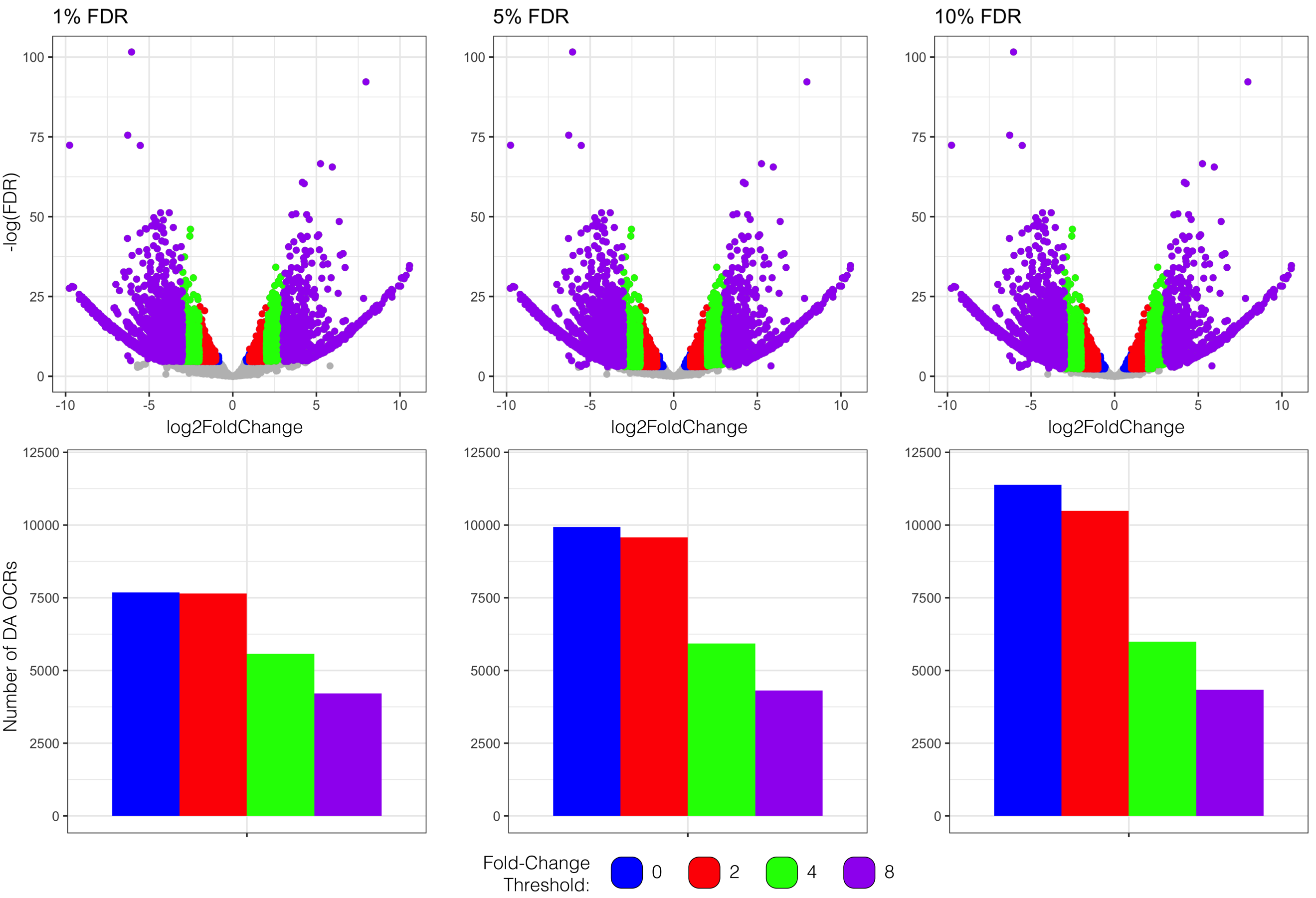

**Supplemental Figure 7:** Number of significantly differentially accessible OCRs between *H. erythrogramma* and *H. tuberculata* at gastrula stage based on different FDR and fold-change cutoff thresholds. Combinations of 1%, 5%, or 10% FDR with absolute fold-change in accessibility of 0, 2, 4, and 8 between species were tested.
